# *In silico* analyses of penicillin binding proteins in *Burkholderia pseudomallei* uncovers SNPs with utility for phylogeography, species differentiation, and sequence typing

**DOI:** 10.1101/2021.10.08.463618

**Authors:** Heather P. McLaughlin, Christopher A. Gulvik, David Sue

**Affiliations:** Biodefense Research and Development Laboratory, Division of Preparedness and Emerging Infections, National Center for Emerging and Zoonotic Infectious Diseases, Centers for Disease Control and Prevention, Atlanta, GA, USA; Zoonoses and Select Agent Laboratory, Division of High-Consequence Pathogens and Pathology, National Center for Emerging and Zoonotic Infectious Diseases, Centers for Disease Control and Prevention, Atlanta, GA, USA

## Abstract

**Background:** *Burkholderia pseudomallei* causes melioidosis. Sequence typing this pathogen can reveal geographical origin and uncover epidemiological associations. Here, we describe *B. pseudomallei* genes encoding putative penicillin binding proteins (PBPs) and investigate their utility for determining phylogeography and differentiating closely related species.

**Methodology & Principal Findings:** We performed *in silico* analysis to characterize 10 PBP homologs in *B. pseudomallei* 1026b. As PBP active site mutations can confer β-lactam resistance in Gram-negative bacteria, PBP sequences in two resistant *B. pseudomallei* strains were examined for similar alterations. Sequence alignments revealed single amino acid polymorphisms (SAAPs) unique to the multidrug resistant strain Bp1651 in the transpeptidase domains of two PBPs, but not directly within the active sites. Using BLASTn analyses of complete assembled genomes in the NCBI database, we determined genes encoding PBPs were conserved among *B. pseudomallei* (n=101) and *Burkholderia mallei* (n=26) strains. Within these genes, single nucleotide polymorphisms (SNPs) useful for predicting geographic origin of *B. pseudomallei* were uncovered. SNPs unique to *B. mallei* were also identified. Based on 11 SNPs identified in two genes encoding predicted PBP-3s, a dual-locus sequence typing (DLST) scheme was developed. The robustness of this typing scheme was assessed using 1,523 RefSeq genomes from *B. pseudomallei* (n=1,442) and *B. mallei* (n=81) strains, resulting in 32 sequence types (STs). Compared to multi-locus sequence typing (MLST), the DLST scheme demonstrated less resolution to support the continental separation of Australian *B. pseudomallei* strains. However, several STs were unique to strains originating from a specific country or region. The phylogeography of Western Hemisphere *B. pseudomallei* strains was more highly resolved by DLST compared to internal transcribed spacer (ITS) typing, and all *B. mallei* strains formed a single ST.

**Significance:** Conserved genes encoding PBPs in *B. pseudomallei* are useful for strain typing, can enhance predictions of geographic origin, and differentiate strains of closely related *Burkholderia* species.

**Author Summary:** *Burkholderia pseudomallei* causes the life-threatening disease melioidosis and is considered a biological threat and select agent by the United States government. This soil-dwelling bacterium is commonly found in regions of southeast Asia and northern Australia, but it is also detected in other tropical and sub-tropical areas around the world. With a predicted global burden of 165,000 annual cases and mortality rate that can exceed 40% without prompt and appropriate antibiotic treatment, understanding the epidemiology of melioidosis and mechanisms of antibiotic resistance in *B. pseudomallei* can benefit public health and safety. Recently, we identified ten conserved genes encoding putative penicillin binding proteins (PBPs) in *B. pseudomallei*. Here, we examined *B. pseudomallei* PBP sequences for amino acid mutations that may contribute to β-lactam resistance. We also uncovered nucleotide mutations with utility to predict the geographical origin of *B. pseudomallei* strains and to differentiate closely related *Burkholderia* species. Based on 11 informative single nucleotide polymorphisms in two genes each encoding a PBP-3, we developed a simple, targeted dual-locus typing approach.

## Introduction

Melioidosis is an emerging but neglected infectious disease caused by the environmental Gram-negative bacterium *Burkholderia pseudomallei*. This soil- and surface water-dwelling microorganism is commonly found in tropical and subtropical regions of Southeast Asia and northern Australia, but also reported in other regions of the Western Hemisphere (WH) [1, 2]. In August 2021, the US Centers for Disease Control and Prevention issued a Health Alert describing a multistate (GA, KS, MN, TX) investigation of non-travel associated melioidosis in four patients [3]. Naturally-acquired melioidosis infections can occur in humans and a wide range of other animals as a result of percutaneous inoculation, inhalation, or ingestion of *B. pseudomallei* [4]. In 2016, a global modeling study predicted that ∼165,000 human melioidosis cases occur annually, and an estimated 89,000 result in death [1]. Infrequent laboratory diagnosis and clinical recognition due to unfamiliarity with the disease and the intrinsic resistance of *B. pseudomallei* to numerous antibiotics can cause delays in treatment leading to poor patient outcomes and mortality rates of up to 50% [5, 6]. *B. pseudomallei* is also among a small group of high-consequence pathogens and toxins regulated in the US in which their misuse could pose a serious threat to public health and safety [7].

Clinically relevant β-lactams used to treat human melioidosis include the cephalosporin ceftazidime, the carbapenems meropenem and imipenem, and the β-lactam/ β-lactam inhibitor combination drug amoxicillin-clavulanic acid [8]. Mechanisms of resistance to these antibiotics have been described in *B. pseudomallei* and involve inactivation of β-lactams as well as modification of β-lactam targets. Mutations in *penA*, or its promoter region that results in overexpression of a β-lactamase, confers resistance to ceftazidime, imipenem, and amoxicillin-clavulanic acid [9-11]. For instance, *B. pseudomallei* isolated from a melioidosis patient in Thailand who succumbed to infection, developed resistance *in vivo* during ceftazidime treatment due to a PenA mutation (Pro167Ser) [12]. Acquired resistance to ceftazidime was also reported due to a reversible gene duplication and amplification event of the genomic region containing *penA* [13]. Understanding and rapidly identifying the mechanisms that contribute to β-lactam resistance in *B. pseudomallei* could inform treatment strategies and improve melioidosis patient outcomes.

Penicillin binding proteins (PBPs) are involved in the final stages of cell wall peptidoglycan synthesis and are conserved among bacteria, with several usually found per species. These proteins determine bacterial cell shape by regulating the localization, timing, and architecture of peptidoglycan polymerization [14]. PBPs are also well-known targets for β-lactams. The covalent binding of a β-lactam antibiotic to the catalytic serine residue at the PBP active site inactivates protein function resulting in inhibition of cell wall synthesis and cell lysis [15, 16]. In bacteria such as *Salmonella enterica, Streptococcus pneumoniae*, and *Helicobacter pylori*, mutations in PBPs within or near conserved active site motifs can result in β-lactam antibiotic resistance by reducing antibiotic binding affinity [17-19]. Chantratita *et al*. demonstrated the loss of PBP-3 in *B. pseudomallei* resulted in ceftazidime resistance [20]. Except for PBP-3, very little is known about PBPs in *B. pseudomallei*. Recently, our group used an *in silico* approach to identify a suite of genes encoding 10 putative PBPs in the *B. pseudomallei* genome [21].

Sequence variations within stable genetic markers in *B. pseudomallei* can be used for: species identification, differentiation of closely related species, characterization of isolates, and also phylogenetic and epidemiological investigations. *Burkholderia mallei* is considered a host-adapted deletion clone of *B. pseudomallei*. As the genes retained by *B. mallei* share ∼99.5% nucleotide identity to corresponding homologs in *B. pseudomallei* [22], the accurate differentiation of these species using molecular-based laboratory tools is difficult but valuable for clinical applications. For example, 16S rRNA gene sequencing rapidly identifies *B. pseudomallei* based on a single nucleotide difference that can reliably discriminate it from *B. mallei* [23]. Polymorphisms within the 16S-23S ribosomal DNA internal transcribed spacer (ITS) have been used to investigate phylogenetic relationships within *B. pseudomallei* and among near-neighbor species [24]. ITS types C, E and CE represented the most endemic *B. pseudomallei* isolates and all isolates of the relative species *B. thailandensis* possessed ITS type A. The ITS allele of *B. mallei* appears monomorphic since all strains were found to have ITS type C [24]. A multi-locus sequence typing (MLST) scheme was also developed for *B. pseudomallei* and closely related species. This molecular typing method is based on sequence variations within seven conserved, housekeeping genes on chromosome I, the larger of its two replicons. MLST demonstrated utility for epidemiological studies and confirmed that *B. mallei* is a clone of *B. pseudomallei*, while the species *Burkholderia thailandesis* is distinct [25].

Despite having a highly recombinant genome, a strong geographic signal is encoded within *B. pseudomallei* and phylogeographic reconstruction of this population is possible [26]. Several factors have led to distinct Asian and Australasian *B. pseudomallei* populations that undergo regional evolution. These factors include the primary mode of transmission (via direct contact with contaminated environments), extremely rare human-to-human transmission, and substantial geographic barriers that restrict gene flow between populations [26]. While a large-scale comparative genomics approach is essential to determine fine-scale population structure and to confirm the true geographic origin of *B. pseudomallei* isolates [27-30], lower resolution typing methods such as ITS and MLST are useful tools for linking melioidosis cases to particular regions. Of the five ITS types exclusive to *B. pseudomallei*, type G was rare in Australia and Southeast Asia, and based on a small number of strains, this type was overrepresented for isolates originating from Africa and the Americas [24]. Testing of additional Western Hemisphere strains confirmed ITS type G was predominant and supported the original hypothesis that a genetic bottle neck took place during dispersal of *B. pseudomallei* to geographic locations outside endemic regions [24, 31]. MLST can be used to define geographical segregation of *B. pseudomallei* by continent and provides a clear distinction between populations originating from Australia and Thailand [32, 33]. However, occasional examples of ST homoplasy have been reported for isolates from different continents that are not actually related [34].

Despite the heavy disease burden and high mortality rate associated with melioidosis, even with aggressive antibiotic treatment [35], melioidosis is not included on the World Health Organization list of neglected tropical diseases and global strategies to address prevention and control are still needed. As the signs and symptoms of melioidosis frequently mimic other diseases, clinical or laboratory diagnosis can be challenging. Prompt diagnosis of this disease as well as timely treatment with appropriate antibiotics are crucial for positive patient outcomes. The genomes of *B. pseudomallei, B. mallei*, and *B. thailandensis* submitted by scientists from across the world to public databases could reveal important markers useful for speciation, predicting antibiotic resistance, phylogeny, or geographic origin.

Here, we utilized an *in silico* approach to characterize PBPs in *B. pseudomallei* and determined their conservation among *B. pseudomallei* isolates as well as closely related species, *B. mallei* and *B. thailandensis*. We also analyzed *B. pseudomallei* PBP sequences for i) amino acid mutations that may confer resistance to β-lactam antibiotics, ii) single nucleotide polymorphisms (SNPs) with utility for species differentiation, and iii) SNPs to infer phylogeographic origins.

## Methods

### *In silico* characterization of PBP homologs in *B. pseudomallei* 1026b

Ten PBP homologs were identified in *B. pseudomallei* 1026b (**Table 1**) using the UniProtKB database (https://www.uniprot.org/). Conserved protein domains were predicted using the Pfam database (http://pfam.xfam.org/) and theoretical molecular weight was calculated using ExPASy (https://web.expasy.org/compute_pi/). NCBI’s Protein BLAST was utilized to find the nearest PBP homologs in *Pseudomonas aeruginosa* PAO1 (taxid:208964) and *Escherichia coli* K-12 (taxid:83333). The nearest homologs produced the most significant alignments to *B. pseudomallei* 1026b PBPs with the lowest Expect (E)-value (NCBI, [36]). Geneious (v.11.1.4) was used to analyze PBP sequences and identify putative enzyme active sites (SXXK, SXN, and KS/TG).

**Table 1.**
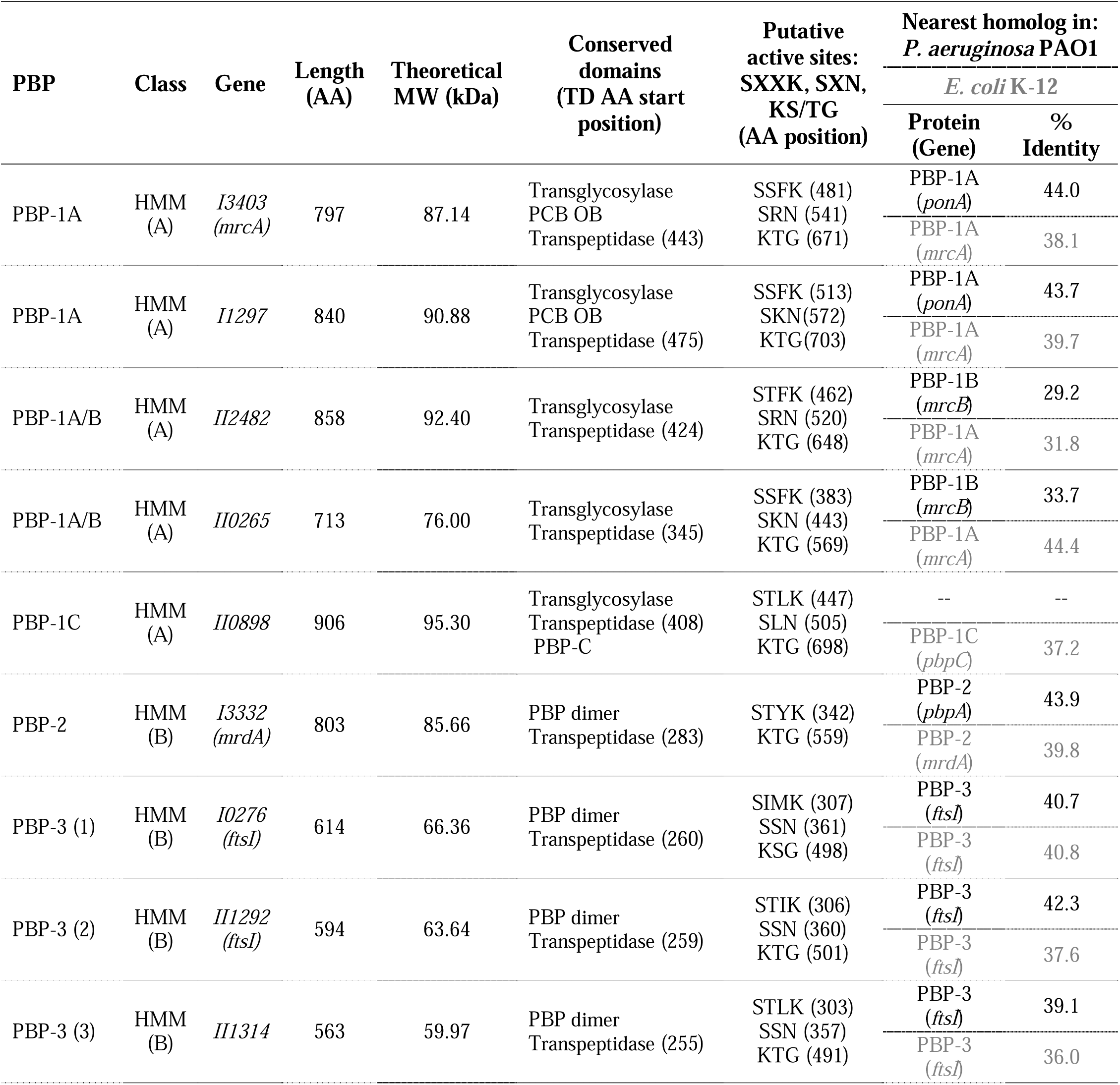
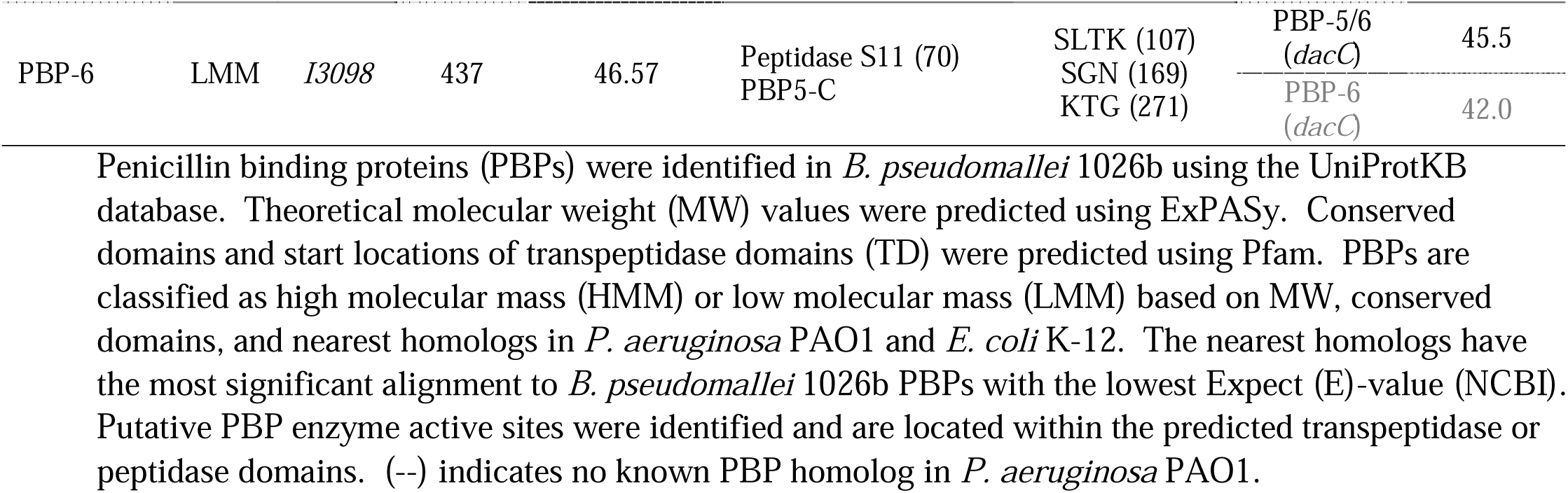
PBP homologs in *B. pseudomallei* 1026b.

### Identification of PBP homologs in genomes of an initial set of *Burkholderia* strains

The nucleotide sequences for the 10 genes encoding PBPs in *B. pseudomallei* 1026b were used as queries for BLASTn analysis. The search set included organisms *B. pseudomallei* (taxid:28450), *B. mallei* (taxid:13373) and *B. thailandensis* (taxid:57975). Default algorithm parameters were selected, with the exception of “max target sequences” which was set to 5,000. This initial set of 144 complete, assembled genomes, including 101 *B. pseudomallei*, 26 *B. mallei*, and 17 *B. thailandensis*, was examined to identify corresponding PBP homologs. These strains, along with their origin, epidemiological information, and NCBI accession numbers, are listed in **Table S1**. Strain typing (MLST, ITS and whole-genome SNP typing) and phylogeography of *B. pseudomallei* from the Western Hemisphere was previously performed and reported by Gee *et. al* [37].

### Analysis of SAAPs in PBP transpeptidase domains

The Pfam database was used to predict the transpeptidase domain (TD) location within each of the 10 PBP homologs in the *B. pseudomallei* 1026b reference strain. Gene sequences obtained from each BLASTn result for 101 *B. pseudomallei* strains were aligned, mapped to the reference strain, translated, and analyzed for single amino acid polymorphisms (SAAPs) within the predicted TDs using Geneious (v11.1.4). The amino acid position of SAAPs and the location of the putative enzyme active sites identified within the TDs are based on sequence alignment to the 1026b reference strain. The Protein Variation Effect Analyzer (PROVEAN) tool (v1.1) [38] was used to predict whether a SAAP affects protein function based on a generated PROVEAN score. A SAAP is predicted to have a ‘deleterious’ effect if the PROVEAN score is ≤ the predefined threshold of -2.5. A SAAP is predicted to have a ‘neutral’ effect if the score is greater than - 2.5.

### Identification and selection of SNPs for the DLST scheme

The nucleotide sequences of *B. pseudomallei* 1026b genes *I0276* and *II1314*, encoding PBP-3 (1) and PBP-3 (3), were used as queries for NCBI’s BLASTn analysis. Gene sequences obtained from BLASTn results for the initial set of 144 *Burkholderia* strains were aligned, mapped to the reference strain, *B. pseudomallei* 1026b, and analyzed for SNPs using Geneious (v11.1.4). Nine SNPs with utility to predict the geographic origin of *B. pseudomallei*, plus two SNPs useful for differentiating *Burkholderia* species (*B. pseudomallei, B. mallei*, and *B. thailandensis*) were identified and selected for DLST. The nucleotide positions described for the 11 DLST SNPs are based on alignment to the 1026b reference strain.

### DLST performance for an expansive set of genomes

All RefSeq genomes of *B. pseudomallei* (n=1525) and *B. mallei* (n=83) were collected from NCBI on Oct. 28^th^, 2019. All *B. mallei* assemblies (n=83) and *B. pseudomallei* assemblies (n=1446) with geographic information deposited in the NCBI BioSample database were evaluated for the presence of genes *I0276* and *II1314* with BLASTn v2.9.0+, using *B. pseudomallei* 1026b as a reference for each sequence. For both genes, the best alignment for each assembly, based on bitscore, was evaluated. A multiple record FastA file (GNU Awk, v4.1.4) was generated using aligned sequences saved from each BLASTn result. Both gene sequence sets were aligned using MUSCLE (v3.8.1551) [39] and visualized in ClustalX [40] to confirm the accuracy of the alignments. Of the 1446 *B. pseudomallei II1314* sequences, 1442 shared >98% nucleotide identity to the 1026b reference strain and were included in the final number of assemblies evaluated in this study. The four remaining sequences in the *II1314* alignment (*B. pseudomallei* strains 3001161896, A193, BP-6260, and BURK081) contained excessive SNPs and gaps, and were excluded based on poor alignment and low nucleotide identity (67%). *I0276* sequences required no further filtering. The BioPython (v1.70) [41] library was used to extract nucleotide data at the 11 positions from both multiple record FastA files and SeqKit (v0.11.0) concatenated the two gene loci for subsequent DLST analysis. The discriminatory power (*D*) of the DLST scheme was calculated using the calculator (http://insilico.ehu.es/mini_tools/discriminatory_power/), where *D* is expressed by the formula of Simpson’s index of diversity [42].

### Compilation of figures

Illustrations of the DLST SNP locations in *B. pseudomallei* 1026b and the DLST SNP-based phylogeographic tree for the initial set of *B. pseudomallei* and *B. mallei* strains (**Fig. 1 and 2**) were generated in Microsoft PowerPoint® for Microsoft 365 MSO (16.0.13801.20840) 64-bit. The more expansive phylogeographic tree (**Fig. 3**), including 1442 *B. pseudomallei* strains resulting in 31 DLSTs, was generated using MPBoot (v1.1.0) [43]. The tree was visualized using the iTol webserver [44] and pie charts were created with the ggplot2 (v3.2.1) library in R (v3.4.4), then edited using InkScape (v0.92.4).

**Figure 1.**
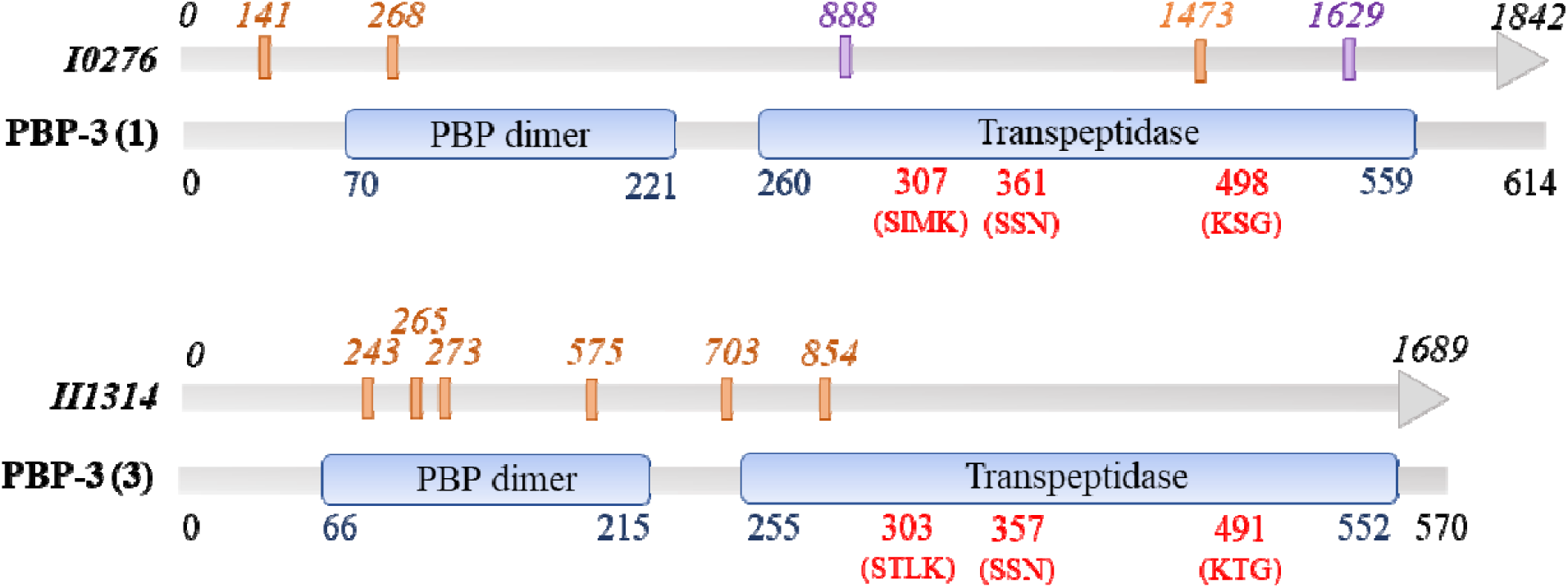
DLST scheme based on 11 nucleotides in two genes (*I0276* and *II1314*) encoding putative PBP-3s. Nucleotide positions are based on sequence alignment to the *B. pseudomallei* 1026b reference strain. Nucleotides with phylogeographic utility (orange font) and utility for differentiating closely related *Burkholderia* species (purple font) are shown. PBP conserved domains (blue bubbles), amino acid position of domains (blue font), active site residues and positions (red font).

**Figure 2.**
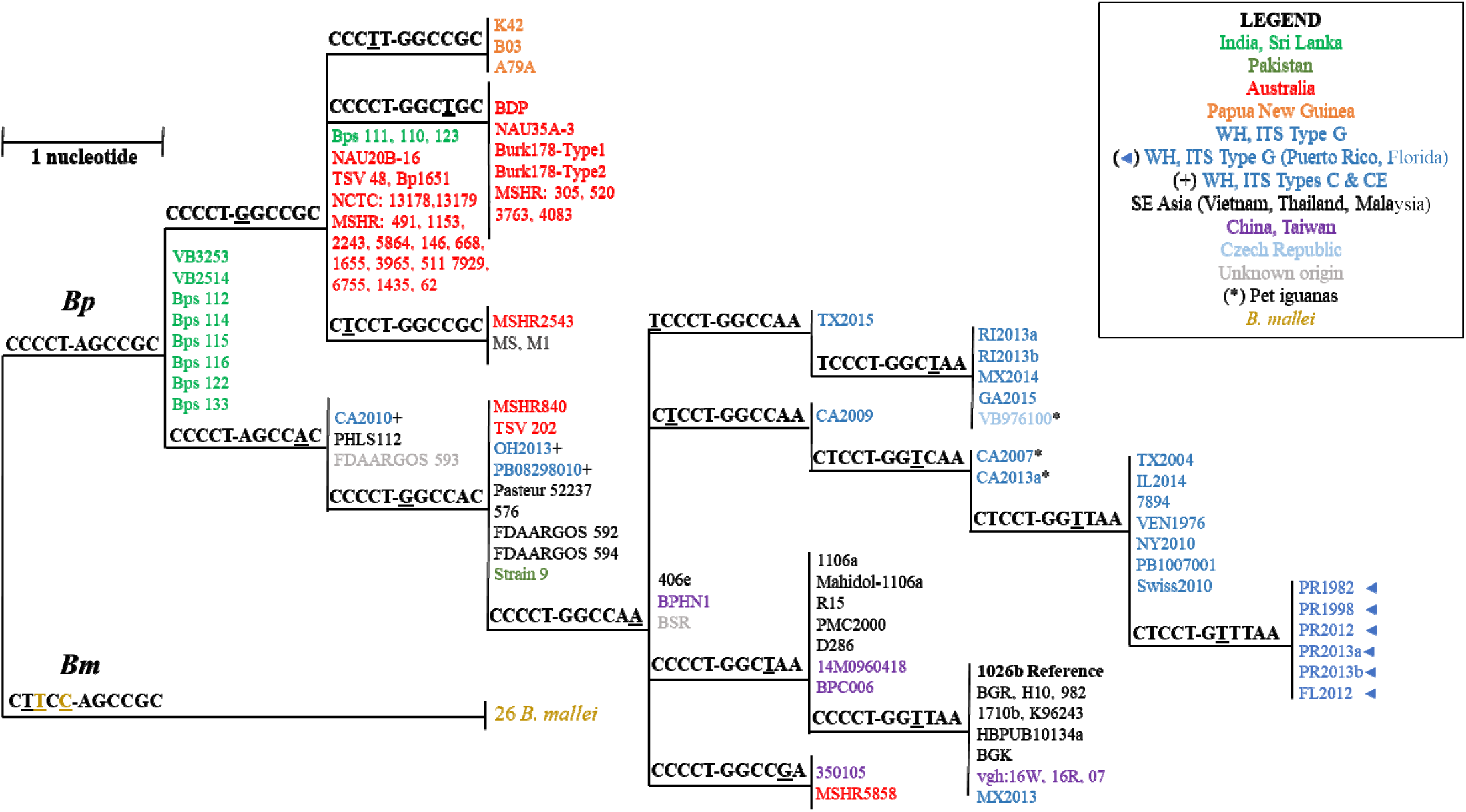
DLST SNP-based phylogeographic tree for the initial set of *B. pseudomallei* (n=101) and *B. mallei* (n=26). Each branch represents isolates with a distinct 11-nucleotide SNP signature determined by DLST. *B. pseudomallei* strains are color-coded by geographic origin and SNPs used to differentiate *B. mallei* strains are shown in dark yellow.

**Figure 3.**
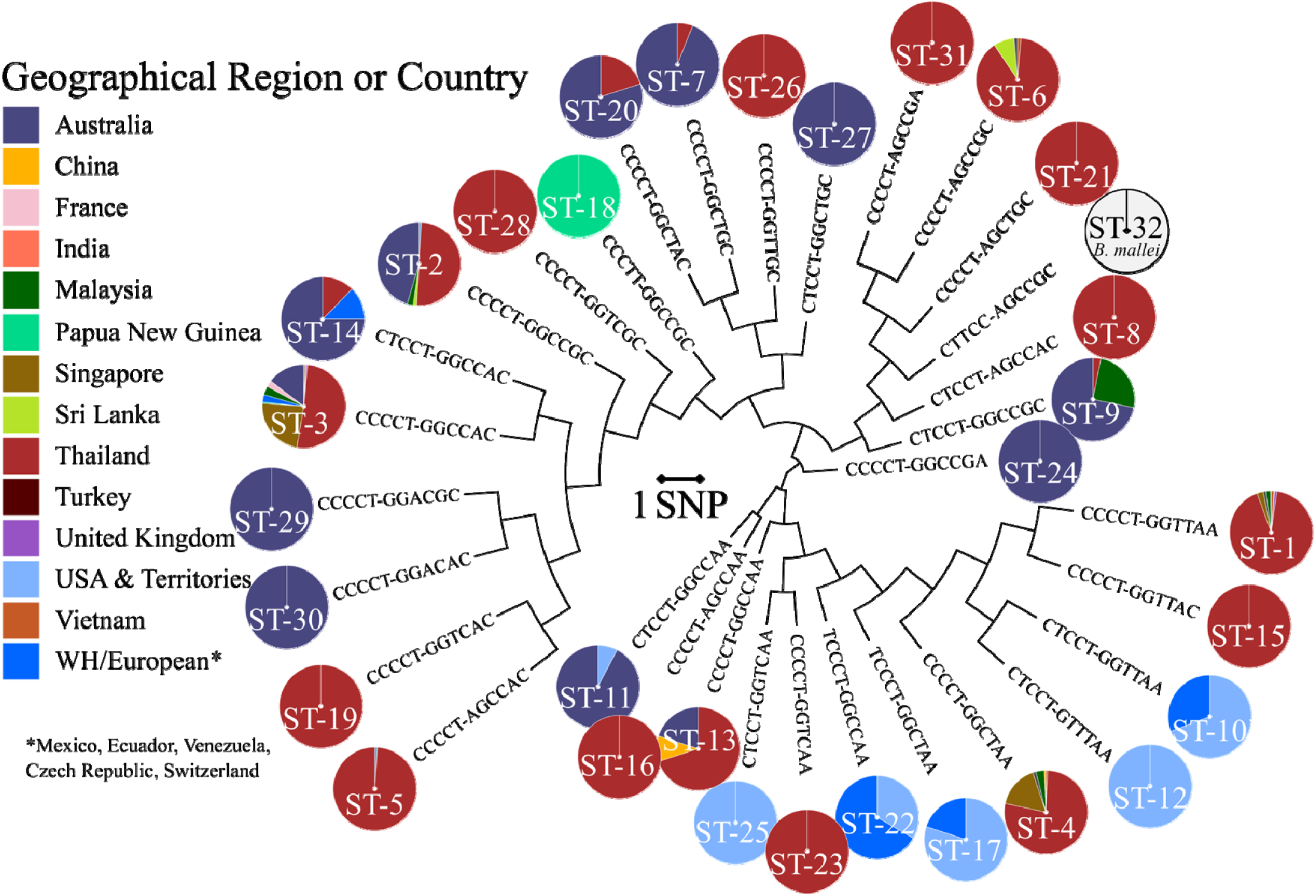
DLST SNP-based phylogeographic tree for the expansive set of *B. pseudomallei* (n=1,442, sequence types 1 to 31) and *B. mallei* (n=81, sequence type 32). *B. pseudomallei* strains are color-coded by geographic origin. Western Hemisphere (WH), asterisk indicates WH and European countries Mexico, Ecuador, Venezuela, Czech Republic, and Switzerland.

## Results/Discussion

### *In silico* characterization of predicted PBPs in *B. pseudomallei* 1026b

Bacterial species have distinctive suites of PBPs and variation exists in the number and redundancy of PBP homologs [45]. Four PBPs are encoded in the *H. pylori* genome, whereas eight and 12 PBPs have been identified in *P. aeruginosa* and *E. coli*, respectively [46-48]. Using the UniProtKB database, we previously identified 10 genes encoding putative PBPs in the *B. pseudomallei* reference strain 1026b [21]. Based on theoretical molecular weight (MW), conserved domains, and the nearest homologs in *P. aeruginosa* and *E. coli, B. pseudomallei* PBPs were classified as high molecular mass (HMM) or low molecular mass (LMM) (**Table 1**). Five HMM, Class-A PBP-1 homologs were identified in *B. pseudomallei* 1026b, each containing a transglycosylase and transpeptidase conserved domain. These proteins range from 713 to 906 amino acids in length with MWs of 95.3 to 76.0 kDa. The MWs of PBP-1 proteins for numerous Gram-negative species have been reported, ranging from 77 to 118 kDa [45]. Both genes encoding PBP-1A homologs (*I3403* and *I1297*) are located on chromosome 1 of the *B. pseudomallei* 1026b genome and share 38.1 to 44.0 % identity to PBP-1As in *P. aeruginosa* PAO1 and *E. coli* K-12, respectively. The predicted PBP-1C, encoded by *II0898* on chromosome 2, was the largest HMM, Class-A protein, and shared 37.2 % identity with PBP-1C in *E. coli* K-12.

Four HMM, Class-B PBP homologs (one PBP-2 and three PBP-3s) were also identified in *B. pseudomallei* 1026b, each containing a transpeptidase conserved domain. The largest of the four, PBP-2, encoded by *I3332* on chromosome 1, was 803 amino acids in length and had a MW of 85.66 kDa. The MWs calculated for the three PBP-3 homologs in *B. pseudomallei* 1026b, encoded by genes *I0276, II1292* and *II1314*, ranged from 59.97 to 66.36 kDa and are comparable to 66 kDa reported previously for PBP-3 in *E. coli* [45]. Protein BLAST analyses revealed *B. pseudomallei* Class-B PBPs share 36.0 to 43.9% identity with the corresponding PBPs in *P. aeruginosa* and *E. coli*. Comparable sequence identity (42%) is reported between PBP-3 homologs of *P. aeruginosa* and *E. coli* [49]. One putative LMM, Class-C PBP-6, encoded by *I3098*, was found on chromosome 1 of *B. pseudomallei* 1026b. This protein represents the smallest of the 10 predicted *B. pseudomallei* PBPs and shares >40% identity to PBP-5/6 and PBP-6 in *P. aeruginosa* PAO1 and *E. coli* K-12, respectively.

The three conserved PBP sequence motifs that form that catalytic center of the active site (SXXK, SXN, and KS/TG) were identified in the transpeptidase domains for nine of the 10 PBP homologs in *B. pseudomallei* 1026b (**Table 1**). For each of these PBPs, the SXXK motifs were located between 54 and 62 residues upstream of the SXN motifs. Corresponding motifs in PBP-3 of *P. aeruginosa* PAO1 are similarly positioned, 55 residues apart [49]. For seven of the nine PBPs, the distance between the SXN and KS/TG motifs was ∼130 residues. This is comparable to the 135 residues that separate the SSN motif from KSG in PBP-3 of *P. aeruginosa* PAO1 [49]. Only two of three active site motifs were found in the *B. pseudomallei* 1026b PBP-2, an STYK tetrad at site 342 and a KTG triad at position 559. It is unclear whether this predicted PBP-2 is functional despite missing the SXN active site sequence, as Tomberg *et al*. [50] demonstrated that an interaction involving the middle residue of this motif was necessary for the transpeptidase function, but not β-lactam binding, of PBP-2 in *Neisseria gonorrhoeae*.

### Identification of PBP homologs among an initial set of *Burkholderia* strains analyzed

The nucleotide sequences for the 10 genes encoding PBPs in *B. pseudomallei* 1026b were used as queries for BLASTn analyses of three *Burkholderia* spp.; *B. pseudomallei* (taxid:28450), *B. mallei* (taxid:13373) and *B. thailandensis* (taxid:57975). An initial set of 143 publicly available, assembled genomes, including 100 additional *B. pseudomallei*, 26 *B. mallei*, and 17 *B. thailandensis* strains were examined to identify genes encoding corresponding PBP homologs. NCBI accession numbers and epidemiological information for these strains can be found in **Table S1**. PBPs in *B. pseudomallei* were more homologous to PBPs in *B. mallei* compared to *B. thailandensis*. With 100% query coverage, BLASTn analysis revealed genes encoding PBPs in *B. mallei* were ≥99% identical to those in the *B. pseudomallei* 1026b genome. Gene sequences of predicted PBPs in *B. thailandensis* were ∼95% identical with query coverages ranging from 85 to 100%.

All ten predicted PBPs identified in *B. pseudomallei* 1026b were conserved in the entire set of *B. pseudomallei* strains. Eight of 10 PBP homologs were conserved in all *B. mallei* genomes evaluated, and the remaining two homologs were present in 25 of 26 genomes. The two exceptions included: *B. mallei* 2002734306 missing one PBP-1A/B, and *B. mallei* SAVP1 missing PBP-1C. Genes encoding 9 of the 10 PBP homologs were present in all *B. thailandensis* genomes examined. However, the third PBP-3 homolog, encoded by *II1314* in *B. pseudomallei* 1026b, was only present in 5 of the 17 *B. thailandensis* strains. Unique PBP profiles analyzed by SDS-PAGE have been used to distinguish species within the *Enterococcus* genus [51]. However, this type of analysis would not prove useful for differentiating *B. pseudomallei* from *B. mallei* and *B. thailandensis*, as several strains from each *Burkholderia* species possess an identical suite of predicted PBPs.

### Examination of PBP transpeptidase domains for mutations in *B. pseudomallei* strains

Alterations in PBPs of Gram-negative bacteria can confer resistance to β-lactams by lowering the affinity of the antibiotic to the active site [17-19]. In this work, we examined the predicted transpeptidase domains (TDs) in *B. pseudomallei* PBPs for single amino acid polymorphisms (SAAPs) in or near active site sequence motifs. The Pfam database was used to predict conserved PBP TDs in *B. pseudomallei* reference strain 1026b. In the PBP-6 and five PBP-1 homologs, the TDs started 37 to 39 residues upstream of the predicted active-site serine residue in the first motif (SXXK). For the four HMM Class-B protein homologs, TDs were positioned ∼47 and 59 residues upstream of the SXXK motif in the three PBP-3s and in PBP-2, respectively (**Table 1**).

The amino acid sequences of the ten putative PBP homologs in the set of 100 *B. pseudomallei* strains were aligned to the 1026b reference strain and examined for alterations. Two strains included in this set have known resistance to β-lactam antibiotics. Based on minimal inhibitory concentration interpretative criteria established by the Clinical Laboratory and Standards Institute, *B. pseudomallei* Bp1651 is considered resistant to amoxicillin-clavulanic acid (AMC), imipenem (IPM), and ceftazidime (CAZ), and *B. pseudomallei* MSHR1655 is resistant to AMC [52]. The reference strain *B. pseudomallei* 1026b is susceptible to all three β-lactams (AMC, IPM, CAZ). No SAAPs were identified directly within any of the three predicted active site motifs for any of the 101 *B. pseudomallei* strains analyzed. In addition, no SAAPs specific to the AMC-resistant strain MSHR1655 were found within the TD domains of the ten PBP homologs.

Antimicrobial resistance markers, including point mutations in the class A β-lactamase encoding *penA* gene, have been described for *B. pseudomallei* Bp1651 [9]. Here, analysis of the TDs in the PBP homologs of this multi-drug resistant strain revealed unique SAAPs that were not present in the other 100 *B. pseudomallei* strains evaluated (**Table 2**). To predict whether these SAAPs would affect protein function, the PROVEAN tool was used to score each mutation individually. The predicted effect was deleterious if the score was ≤ -2.5 and neutral if the score was > -2.5. Two potentially deleterious SAAPs exclusive to strain Bp1651 were found: G608D, located 49 residues downstream of the KTG active site motif in the PBP-2 homolog, and G530R, located 25 residues downstream of the SLN motif in the PBP-1C homolog. A neutral effect on protein function was calculated for the second amino acid substitution in the PBP-1C homolog at position 627. One additional predicted deleterious SAAP (G495S), 25 residues upstream of the second active site motif in a PBP-1A/B homolog, was found in *B. pseudomallei* strains Bp1651 and MSHR1153. The functional contributions of these SAAPs in Bp1651 to β-lactam resistance remain unknown.

**Table 2.** Single amino acid polymorphisms found in PBP transpeptidase domains of *B. pseudomallei* Bp1651.

| PBP Homolog | Corresponding Gene in Bp1651 | SAAP in TD (AA position) | Location relative to putative active site | PROVEAN |  |
| --- | --- | --- | --- | --- | --- |
|  |  |  |  | Score | Predicted Affect |
| PBP-2 | *TR70_2681* | G ► D (608) | (+) 49 AA, KTG | - 4.094 | Deleterious |
| PBP-1A/B | *TR70_5295* | G ► S (495)* | (-) 25 AA, SRN | -4.091 | Deleterious |
| PBP-1C | *TR70_6206* | G ► R (530) | (+) 25 AA, SLN | -7.500 | Deleterious |
|  |  | E ► Q (627) | (-) 72 AA, KTG | -0.238 | Neutral |

The amino acid (AA) position of single amino acid polymorphisms (SAAPs) in predicted PBP transpeptidase domains (TD) of multi-drug resistant Bp1651 strain are based on sequence alignment to the 1026b reference strain. Location relative to the nearest active site, (-) upstream or (+) downstream. SAAPs listed are unique to Bp1651 and not found in the other 100 *B. pseudomallei* strains analyzed. (\*) SAAP shared only with *B.* *pseudomallei* MSHR1153. The Protein Variation Effect Analyzer (PROVEAN) tool was used to predict whether a SAAP affects protein function. The predicted affect is deleterious if the score $\leq -2.5$ and neutral if the score is $>$ $-2.5$ .

### Utility of PBP gene SNPs for differentiation of closely related *Burkholderia* species

As a result of the high degree of phenotypic and genotypic overlap between *B. pseudomallei, B. mallei*, and *B. thailandensis*, simple molecular approaches to differentiate these species are important for epidemiological studies and clinical applications. Some PCR-based methodologies targeting open reading frames including *16S rRNA, bimA*, and *fliC* genes, have been described for the *Burkholderia pseudomallei* complex and were summarized by Lowe *et al*. [53]. As part of this work, gene sequences encoding each of the ten putative PBP homologs in the initial set of *Burkholderia* strains (**Table S1**) were aligned to the *B. pseudomallei* 1026b reference strain and examined for species-specific SNPs. Several PBP gene homologs contained mutations with utility to differentiate the three closely related *Burkholderia* spp. (**Table 3**). The nucleotide positions for these mutations are reported based on sequence alignments to the 1026b reference strain. The combination of two SNPs at positions 888 and 1629 in the gene encoding the PBP-3 (1) homolog was found to be unique to each of the three *Burkholderia* spp. For 100% of the isolates analyzed at these two sites, we observed nucleotides C/T for *B. pseudomallei* (n=101), T/C for *B. mallei* (n=26) and C/C for *B. thailandensis* (n=17).

**Table 3.**
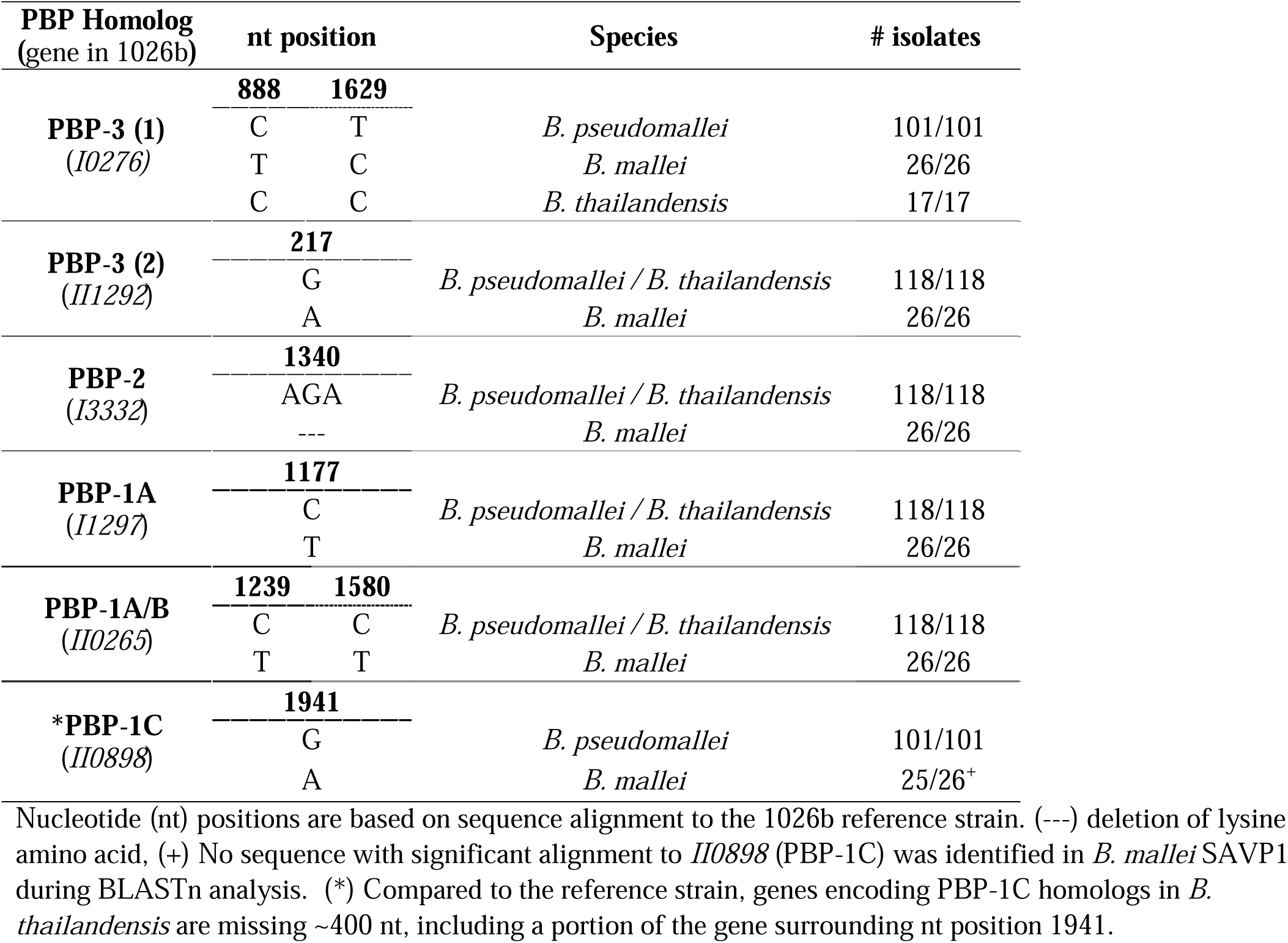
PBP SNPs with utility for differentiation of *Burkholderia* species.

SNPs unique to *B. mallei* were identified in genes encoding five predicted PBPs (**Table 3**). For instance, an adenine at position 217 in PBP-3 (2), a thymine at position 1177 in a PBP-1A, and two thymine nucleotides at positions 1239 and 1580 in a PBP-1A/B could be used to differentiate *B. mallei* from both *B. pseudomallei* and *B. thailandensis*. All *B. mallei* strains (26/26) also contained a three-nucleotide deletion starting at position 1340 in genes encoding the putative PBP-2 homolog. This mutation resulted in the deletion of a lysine for *B. mallei*, which was not observed in any of the 118 *B. pseudomallei* or *B. thailandensis* strains evaluated. Except for *B. mallei* strain SAVP1, which does not contain the gene encoding the predicted PBP-1C, a SNP at position 1941 could be used to differentiate *B. pseudomallei* from *B. mallei*. Compared to the 1026b reference strain, genes encoding PBP-1C homologs in *B. thailandensis* are missing ∼400 nucleotides, including the portion of the gene surrounding the nucleotide at position 1941. Of the 10 predicted PBPs, PBP-3 (1) proved most useful in improving our ability to differentiate *B. pseudomallei, B. mallei*, and *B. thailandensis*.

### Utility of PBP SNPs for predicting geographic origin of *B. pseudomallei*

Phylogeographic reconstruction of *B. pseudomallei* has been demonstrated using both high resolution comparative genomics and lower resolution typing methods such as ITS [24, 27]. To investigate whether isolates could be assigned to a geographic region gene sequences encoding each of the ten putative PBP homologs in 100 *B. pseudomallei* genomes were aligned to the *B. pseudomallei* 1026b reference strain and examined for SNPs useful for phylogeography. In the initial set of 101 *B. pseudomallei* genomes analyzed (**Table S1**), 75 strains originated from the Eastern Hemisphere (EH); 33 strains from Australia and Papua New Guinea, and 42 strains from 8 countries in Asia and Southeast Asia. The remaining 26 *B. pseudomallei* strains were isolated from clinical or environmental samples in the Western Hemisphere (WH); 23 of which have ITS type G, common to strains from the WH, and 3 isolates have ITS type C or CE, supporting an original origin outside the WH, most likely Asia [37].

Several SNPs with phylogeographic utility were identified in genes encoding putative PBPs in the initial set of *B. pseudomallei* strains. For example, analysis of predicted PBP-1A/B gene homologs, aligned to *II0265* in the *B. pseudomallei* 1026b reference strain, revealed a SNP at position 168 shared by all ITS type G, WH strains (23/23). Only 4 strains originating from the EH shared the same SNP. A SNP unique to WH strains originating from Puerto Rico and Florida was also identified in gene sequences encoding PBP-2 homologs at position 108. Another SNP more prevalent to strains originating from Australia and Papua New Guinea (27/33) was identified in gene sequences encoding the second predicted PBP-1A/B homolog found at nucleotide position 2,330. However, the two genes containing the most SNPs with utility to predict geographic origin of *B. pseudomallei* encode the predicted PBP-3 (1) and PBP-3 (3) homologs. Furthermore, as SNPs in the first PBP-3 gene homolog could also be used to differentiate *B. pseudomallei* from *B. mallei* and *B. thailandensis*, these two loci were selected for the Dual-Locus Sequence Typing (DLST) approach.

### Development of a Dual-Locus Sequence Typing approach

DLST is a molecular biology technique that uses unique allelic profiles in two loci to characterize and type bacterial species. Implementation of DLST typing schemes has been demonstrated for bacterial pathogens such as methicillin-resistant *Staphylococcus aureus* and *P. aeruginosa* [54, 55]. Here, a DLST scheme was developed using polymorphic sites at 11 nucleotide positions in two conserved *B. pseudomallei* loci encoding PBP-3 homologs (**Table 4, Fig. 1**). The nucleotide positions reported herein are based on sequence alignment to the *I0276* and *II1314* genes, encoding PBP-3 (1) and PBP-3 (3), respectively, in the *B. pseudomallei* 1026b reference strain. SNPs at 9 of the 11 positions were chosen for their phylogeographic utility, and the other 2 SNPs were useful in differentiating *Burkholderia* spp. The DLST approach was first tested using gene sequences encoding PBP-3 (1) and PBP-3 (3) homologs in the initial set of *B. pseudomallei* (n=101) and *B. mallei* (n=26) strains.

**Table 4.**
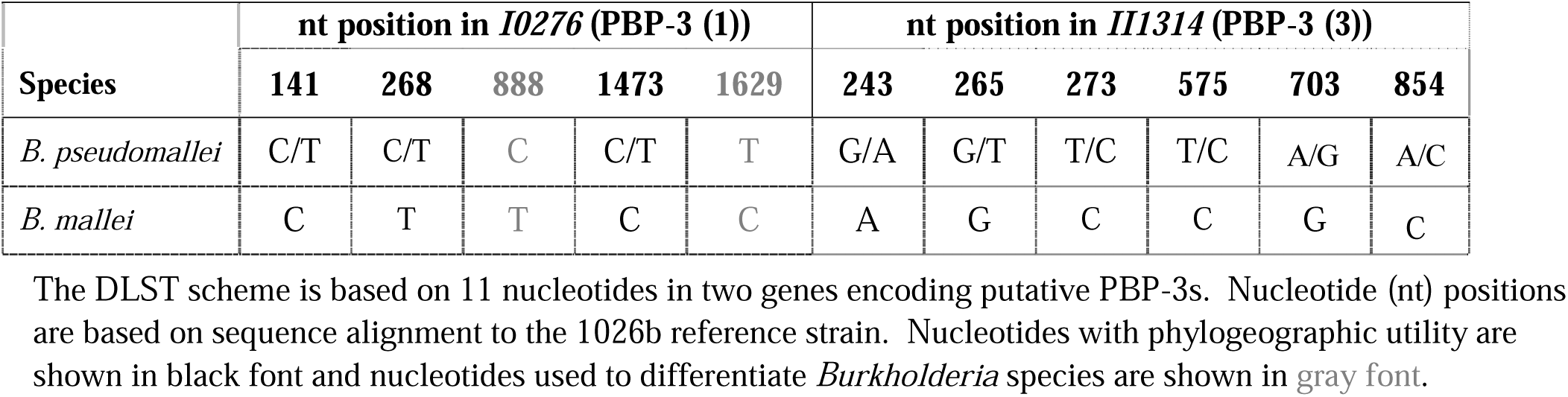
Dual-locus sequence typing scheme.

DLST results for the initial set of 127 *Burkholderia* strains are depicted in a phylogeographic SNP tree (**Fig. 2**) and summarized in **Table S2**. Each branch of the SNP tree represents a group of strains with a distinct 11-nucleotide SNP signature. Directly adjacent branches differ by one SNP (underlined), as indicated by the size bar. In **Fig. 1**, *B. pseudomallei* strains are color-coded by geographic origin and SNPs used to differentiate *B. mallei* strains are shown in dark yellow. All *B. mallei* strains (26/26) shared the same SNP signature (CTTCC-AGCCGC) and clustered together. At the initial bifurcation point of the SNP tree, *B. pseudomallei* strains originating from India and Sri Lanka (in green) group together. Geographically, these countries are in proximity, sharing a maritime border. The other three isolates from Sri Lanka (Bps 111, 110, and 123) share 10 of 11 DLST signature SNPs (**Fig. 2**).

A SNP unique to *B. pseudomallei* strains from Papua New Guinea was identified at nucleotide position 1473 in the first DLST locus, and as a result these three strains formed their own branch on the tree (CCCTT-GGCCGC) (**Fig. 2**). The majority of *B. pseudomallei* strains originating from Australia (26/30), located just south of Papua New Guinea, share 2 DLST signatures and differ from Papua New Guinea strains by only one or two SNPs. MLST has been used to study the origins of isolates from Papua New Guinea [56]; three unique sequence types (STs) were resolved and phylogenetic analysis revealed they were located in clades mainly dominated by isolates of Australian origin. The six DLST signatures in the lower portion of the phylogeographic SNP tree mainly consisted of *B. pseudomallei* strains originating from Southeast Asia (Vietnam, Thailand, and Malaysia), China, and Taiwan (**Fig. 2**). One of which, CCCCT-GGTTAA, contained the most isolates from Thailand (6), including the 1026b reference strain (in bold). Outliers included Australian strains MSHR840 and TSV202 which were assigned to a DLST branch along with five Asian isolates. However, *B. pseudomallei* strains with ITS type C and CE; CA2010, OH2013, and PB08298010 from the WH, and the Australian strain MSHR5858 have all been shown to be more closely related to strains originating in Asia [29, 37, 57]; indeed, these four clinical isolates shared SNP signatures with strains from several Asian countries.

For WH *B. pseudomallei* strains, the DLST system demonstrated higher resolution compared to ITS typing and some DLST groups could be associated with geographic origin. Based on DLST SNP signatures, *B. pseudomallei* ITS Type G isolates from the WH (in blue) could be differentiated into several distinct groups (**Fig. 2**). All WH strains evaluated in this study that originated from Puerto Rico and Florida (6/6) populated a single branch on the SNP tree (CTCCT-GTTTAA). This is consistent with core genome SNP analysis results which showed these six isolates made up a distinct subclade within the WH clade [37]. Moreover, two MLST type 518 *B. pseudomallei* strains, CA2007 and CA2013a, both isolated from pet iguanas in California [37], clustered together using the DLST approach (CTCCT-GGTCAA). The largest DLST group, with SNP signature CTCCT-GGTTAA, consisted of seven WH *B. pseudomallei* strains. One outlier was an ITS Type G isolate from a California patient (MX2013). DLST placed this WH strain in a group with *B. pseudomallei* strains from Southeast Asia and Taiwan. Interestingly, this patient had travel history or residence in Mexico and had prior military service in Vietnam [37] (**Table S1**).

### Testing DLST performance with an expansive set of *B. pseudomallei* and *B. mallei* genomes

The DLST approach was challenged using an extensive set of *B. pseudomallei* and *B. mallei* RefSeq genomes collected from NCBI. All *B. pseudomallei* assemblies (n=1,446) with geographic information deposited in the NCBI BioSample database and *B. mallei* assemblies (n=83) were evaluated for the presence of genes encoding predicted PBP-3 (1) and PBP-3 (3) homologs (*I0276* and *II1314*) with BLASTn, using *B. pseudomallei* 1026b as a reference for each sequence. Both loci were highly conserved in *B. pseudomallei*, each exhibiting >98% nucleotide identity to the reference. Only four of 1,446 *B. pseudomallei* strains were excluded from subsequent DLST analysis based on poor alignment and low nucleotide identity to the second locus, *II1314*. The two putative PBP-3 gene homologs were also highly conserved in *B. mallei*, with only two of 83 strains missing the second locus. As a result, DLST typing performance was ultimately assessed using gene sequences from 1,442 *B. pseudomallei* and 81 *B. mallei* strains. Both gene sequence sets were aligned to the *B. pseudomallei* 1026b reference strain, and nucleotide data at the 11 polymorphic positions were extracted and concatenated for subsequent DLST analysis.

DLST of 1,523 *Burkholderia* strains resulted in 32 sequence types (STs); 31 STs for *B. pseudomallei* strains and one ST (ST-32) for all *B. mallei* strains (**Table S3**). *B. pseudomallei* STs were assigned in numerical order in accordance with the number of strains in each ST, highest to lowest. Based on this typing nomenclature, STs were assigned in retrospect to the initial set of 101 *B. pseudomallei* and 26 *B. mallei* strains used to develop the DLST approach (**Table S2**). To assess the discriminatory power (*D*) of the DLST method, a single numerical index of discrimination [42] was calculated based on the probability that two unrelated, randomly sampled *B. pseudomallei* or *B. mallei* strains from our test population (n=1,523) would be placed in different typing groups. Predicated on 32 STs, the *D* value of this DLST was 0.8512.

DLST data was used to construct a phylogeographic tree (**Fig. 3**). Individual branches depict the unique SNP signature, or allelic profile, for each ST. The geographic origins of strains assigned to each ST are represented by color-coded pie charts at each terminal node. The largest number of *B. pseudomallei* strains (n=451) was assigned to ST-1, of which ∼98% (n=440) geographically originate from Southeast Asia (Thailand, Singapore, Malaysia and Vietnam). No Australian isolates were assigned to ST-1, and the four ST-1 strains described as originating from the United Kingdom are in fact laboratory cultures of the Thai *B. pseudomallei* strain K96243 (**Table S3**). The *B. pseudomallei* 1026b reference strain is among the 416 isolates from Thailand that belong this DLST profile.

DLST revealed only *B. pseudomallei* strains with Southeast and East Asian origin (n=157) belonged to ST-4 (**Table S3**). Nine STs unique to isolates from Thailand (n=63) were also resolved using DLST profiling. These STs are depicted in tree branches with entirely red pie charts (**Fig. 3**). Additionally, a tenth ST (ST-5) included 91 *B. pseudomallei* from Thailand and one strain from the USA; the latter, CA2010, is ITS Type C and is more closely related to *B. pseudomallei* strains from Southeast Asia [37]. ST-18 consisted of five isolates specifically from Papua New Guinea; three from our initial set of *B. pseudomallei* genomes analyzed (strains K42, B03, and A79A) and two additional isolates from this more expansive DLST analysis (**Table S2** and **Table S3**). One other Papua New Guinean *B. pseudomallei* strain (MSHR139) was profiled by DLST and assigned to ST-3. Only the country of isolation is listed in the SAMN02443743 sample information on NCBI, so it is unknown whether this strain was isolated from a melioidosis patient with travel history to other geographic locations. All *B. pseudomallei* strains with French (n=5) and Pakistani (n=3) origin clustered together in ST-3.

While the majority of *B. pseudomallei* isolates separate into 2 phylogenetic groups, Australia and Southeast Asia/rest of the world, a single strain (MSHR5858) with a unique MLST sequence type (ST-562) is present in northern Australia, Taiwan, and southern China [58]. Although we observe four STs specific to a small number of Australian isolates (n=6), this DLST scheme does not completely support the separation of Australasian and Asian *B. pseudomallei* clades. Comparable to ITS typing [24], we observed several STs (7) populated with strains originating from both Thailand and Australia (**Fig. 3** and **Table S3**). Two of these types, ST-7 and ST-9, were more common to Australia, encompassing 94% and 72% of the total number of strains. Twelve of 13 *B. pseudomallei* isolates belonging to ST-11 were from Australia and the other was an outlier, CA2009, an ITS type G strain from the WH. Prior to this more extensive DLST analysis, strain CA2009 resided alone on its own branch in the phylogeographic SNP tree based on the initial set of 101 B. *pseudomallei* strains (**Fig. 2**).

Five STs were observed that were unique to 34 *B. pseudomallei* strains from the WH. ST-10 was the most common DLST profile for WH isolates (n=13) followed by ST-12 (n=11). Consistent with our preliminary DLST analysis which included six *B. pseudomallei* strains isolated from clinical and environmental samples in Puerto Rico and Florida, 6 additional strains from this same geographical region all clustered together in ST-12 (**Fig 2, Fig. 3, Table S3**). DLST sequence type ST-22 contained three WH strains, two originating from Mexico and the other, strain TX2015, isolated from a melioidosis patient with travel history to Mexico [37]. Strains CA2007 and CA2013a, which were both isolated from *Iguana iguana*, remain the only two of 1,442 strains assigned to ST-25. The two WH *B. pseudomallei* strains, designated as American in origin, that fall into ST-2 along with 218 strains from the EH are strain Bp1651, which formerly comes from Australia and 2014002816, which is a clinical isolate from a patient in Maryland with travel history to Africa.

## Conclusions

The true global burden of *B. pseudomallei* infections is likely underestimated due to several factors including difficulty of diagnosis, insufficient methods for conventional identification, and limited diagnostic facilities [59]. Diagnosis and epidemiological analysis of *B. pseudomallei* are critical to ensure positive patient outcomes and investigate outbreaks, however, resource constraints may limit the laboratory techniques employed for routine testing. Rapid, low-cost, and easy to perform methods that produce unambiguous results, portable between laboratories, may be more feasible to implement in such settings.

We characterized genes encoding 10 predicted PBPs that were conserved among sequenced *B. pseudomallei* and *B. mallei* strains. Within these genes, SNPs with utility for phylogeography and species differentiation were uncovered, markedly in those encoding the predicted PBP-3 (1) and PBP-3 (3) homologs. Using 11 polymorphic nucleotides identified within these two loci, a simple DLST typing scheme was developed and challenged with sequence data from over 1500 *B. pseudomallei* and *B. mallei* strains. The willingness of research scientists worldwide to share *B. pseudomallei* and *B. mallei* genome sequences in publicly accessible databases strengthened this work. While WGS offers the most comprehensive (and therefore has the potential to be the most accurate) method for determining the geographic origin of *B. pseudomallei*, lower resolution techniques such as MLST and ITS typing are useful tools for associating melioidosis cases to particular regions. A limitation of the DLST described herein is the reduced discriminatory power compared to WGS and MLST. However, the DLST approach relies on only two gene targets and could be easily operationalized into a PCR test from a culture isolate (or optimized for testing directly from certain clinical specimens). This simple test could be used to rapidly discern strains of closely related *Burkholderia* spp. and perform some phylogeographic reconstruction, most notably for WH *B. pseudomallei* isolates. In summary, sequence typing methods based on conserved genes encoding PBPs in *B. pseudomallei* may be used to improve our current, targeted typing schemes, enhance our ability to link genetic data with geographic origin, and help differentiate closely related *Burkholderia* species, especially in settings where WGS may not be feasible.

## Acknowledgments

We thank Zachary Weiner, Jay Gee, and Mindy Glass Elrod in the Division of High-Consequence Pathogens and Pathology at the Centers for Disease Control and Prevention for their technical expertise and review of this manuscript.

## Disclaimer

The findings and conclusions in this report are those of the authors and do not necessarily represent the official position of the Centers for Disease Control and Prevention. Use of trade names is for identification only and does not imply endorsement by the Centers for Disease Control and Prevention.

## Notes

### Competing Interest Statement

The authors have declared no competing interest.

## References

1. Limmathurotsakul D, Golding N, Dance DA, Messina JP, Pigott DM, Moyes CL, et al. Predicted global distribution of Burkholderia pseudomallei and burden of melioidosis. Nat Microbiol. 2016;1(1):15008.

2. Benoit TJ, Blaney DD, Doker TJ, Gee JE, Elrod MG, Rolim DB, et al. A review of melioidosis cases in the Americas. Am J Trop Med Hyg. 2015;93(6):1134–9.

3. CDC. New Case Identified: Multistate Investigation of Non-travel Associated Burkholderia pseudomallei Infections (Melioidosis) in Four Patients: Georgia, Kansas, Minnesota, and Texas—2021. https://emergency.cdc.gov/han/2021/han00448.asp. Cited September 9, 2021.

4. Limmathurotsakul D, Kanoksil M, Wuthiekanun V, Kitphati R, deStavola B, Day NP, et al. Activities of daily living associated with acquisition of melioidosis in northeast Thailand: a matched case-control study. PLoS Negl Trop Dis. 2013;7(2):e2072.

5. Wiersinga WJ, Virk HS, Torres AG, Currie BJ, Peacock SJ, Dance DA, et al. Melioidosis. J Nature reviews Disease primers. 2018;4:17107.

6. Cheng AC, Currie BJ. Melioidosis: epidemiology, pathophysiology, and management. Clin Microbiol Rev. 2005;18(2):383–416.

7. Program FSA. Select Agent and Toxins List. https://www.cdc.gov/selectagent/SelectAgentsandToxinsList.html. Cited September 9, 2021.

8. Lipsitz R, Garges S, Aurigemma R, Baccam P, Blaney DD, Cheng AC, et al. Workshop on treatment of and postexposure prophylaxis for Burkholderia pseudomallei and B. mallei Infection, 2010. Emerg Infect Dis. 2012;18(12):e2.

9. Bugrysheva JV, Sue D, Gee JE, Elrod MG, Hoffmaster AR, Randall LB, et al. Antibiotic Resistance Markers in Burkholderia pseudomallei Strain Bp1651 Identified by Genome Sequence Analysis. Antimicrob Agents Chemother. 2017;61(6).

10. Rholl DA, Papp-Wallace KM, Tomaras AP, Vasil ML, Bonomo RA, Schweizer HP. Molecular Investigations of PenA-mediated beta-lactam Resistance in Burkholderia pseudomallei. Front Microbiol. 2011;2:139.

11. Sarovich DS, Price EP, Von Schulze AT, Cook JM, Mayo M, Watson LM, et al. Characterization of ceftazidime resistance mechanisms in clinical isolates of Burkholderia pseudomallei from Australia. PLoS One. 2012;7(2):e30789.

12. Sarovich DS, Price EP, Limmathurotsakul D, Cook JM, Von Schulze AT, Wolken SR, et al. Development of ceftazidime resistance in an acute Burkholderia pseudomallei infection. Infect Drug Resist. 2012;5:129–32.

13. Chirakul S, Somprasong N, Norris MH, Wuthiekanun V, Chantratita N, Tuanyok A, et al. Burkholderia pseudomallei acquired ceftazidime resistance due to gene duplication and amplification. Int J Antimicrob Agents. 2019;53(5):582–8.

14. Popham DL, Young KD. Role of penicillin-binding proteins in bacterial cell morphogenesis. Curr Opin Microbiol. 2003;6(6):594–9.

15. Georgopapadakou N, Hammarström S, Strominger JJPotNAoS. Isolation of the penicillin-binding peptide from D-alanine carboxypeptidase of Bacillus subtilis. Proc Natl Acad Sci USA. 1977;74(3):1009–12.

16. Bush K, Bradford PAJCSHpim. β-Lactams and β-lactamase inhibitors: an overview. 2016;6(8):a025247.

17. Sun S, Selmer M, Andersson DI. Resistance to beta-lactam antibiotics conferred by point mutations in penicillin-binding proteins PBP3, PBP4 and PBP6 in Salmonella enterica. PLoS One. 2014;9(5):e97202.

18. Contreras-Martel C, Dahout-Gonzalez C, Martins ADS, Kotnik M, Dessen AJJomb. PBP active site flexibility as the key mechanism for β-lactam resistance in pneumococci. 2009;387(4):899–909.

19. Rimbara E, Noguchi N, Kawai T, Sasatsu M. Mutations in penicillin-binding proteins 1, 2 and 3 are responsible for amoxicillin resistance in Helicobacter pylori. J Antimicrob Chemother. 2008;61(5):995–8.

20. Chantratita N, Rholl DA, Sim B, Wuthiekanun V, Limmathurotsakul D, Amornchai P, et al. Antimicrobial resistance to ceftazidime involving loss of penicillin-binding protein 3 in Burkholderia pseudomallei. Proc Natl Acad Sci U S A. 2011;108(41):17165–70.

21. McLaughlin HP, Bugrysheva J, Sue D. Optical microscopy reveals the dynamic nature of B. pseudomallei morphology during beta-lactam antimicrobial susceptibility testing. BMC Microbiol. 2020;20(1):209.

22. Schell MA, Lipscomb L, DeShazer D. Comparative genomics and an insect model rapidly identify novel virulence genes of Burkholderia mallei. J Bacteriol. 2008;190(7):2306–13.

23. Gee JE, Sacchi CT, Glass MB, D. BK, Weyant RS, Levett PN, et al. Use of 16S rRNA gene sequencing for rapid identification and differentiation of Burkholderia pseudomallei and B. mallei. J Clin Microbiol. 2003;41(10):4647–54.

24. Liguori AP, Warrington SD, Ginther JL, Pearson T, Bowers J, Glass MB, et al. Diversity of 16S-23S rDNA internal transcribed spacer (ITS) reveals phylogenetic relationships in Burkholderia pseudomallei and its near-neighbors. PLoS One. 2011;6(12):e29323.

25. Godoy D, Randle G, Simpson AJ, Aanensen DM, Pitt TL, Kinoshita R, et al. Multilocus sequence typing and evolutionary relationships among the causative agents of melioidosis and glanders, Burkholderia pseudomallei and Burkholderia mallei. J Clin Microbiol. 2003;41(5):2068–79.

26. Pearson T, Giffard P, Beckstrom-Sternberg S, Auerbach R, Hornstra H, Tuanyok A, et al. Phylogeographic reconstruction of a bacterial species with high levels of lateral gene transfer. BMC Biol. 2009;7:78.

27. Chewapreecha C, Holden MT, Vehkala M, Valimaki N, Yang Z, Harris SR, et al. Global and regional dissemination and evolution of Burkholderia pseudomallei. Nat Microbiol. 2017;2:16263.

28. Sarovich DS, Garin B, De Smet B, Kaestli M, Mayo M, Vandamme P, et al. Phylogenomic Analysis Reveals an Asian Origin for African Burkholderia pseudomallei and Further Supports Melioidosis Endemicity in Africa. mSphere. 2016;1(2).

29. Price EP, Sarovich DS, Smith EJ, MacHunter B, Harrington G, Theobald V, et al. Unprecedented Melioidosis Cases in Northern Australia Caused by an Asian Burkholderia pseudomallei Strain Identified by Using Large-Scale Comparative Genomics. Appl Environ Microbiol. 2016;82(3):954–63.

30. Gee JE, Gulvik CA, Castelo-Branco DS, Sidrim JJ, Rocha MF, Cordeiro RA, et al. Genomic Diversity of Burkholderia pseudomallei in Ceara, Brazil. Msphere. 2021;6(1):e01259–20.

31. Gee JE, Allender CJ, Tuanyok A, Elrod MG, Hoffmaster AR. Burkholderia pseudomallei type G in Western Hemisphere. Emerg Infect Dis. 2014;20(4):682–4.

32. Vesaratchavest M, Tumapa S, Day NP, Wuthiekanun V, Chierakul W, Holden MT, et al. Nonrandom distribution of Burkholderia pseudomallei clones in relation to geographical location and virulence. J Clin Microbiol. 2006;44(7):2553–7.

33. Cheng AC, Godoy D, Mayo M, Gal D, Spratt BG, Currie BJ. Isolates of Burkholderia pseudomallei from Northern Australia are distinct by multilocus sequence typing, but strain types do not correlate with clinical presentation. J Clin Microbiol. 2004;42(12):5477–83.

34. De Smet B, Sarovich DS, Price EP, Mayo M, Theobald V, Kham C, et al. Whole-genome sequencing confirms that Burkholderia pseudomallei multilocus sequence types common to both Cambodia and Australia are due to homoplasy. J Clin Microbiol. 2015;53(1):323–6.

35. Dance DA. Melioidosis: the tip of the iceberg? Clin Microbiol Rev. 1991;4(1):52–60.

36. Camacho C, Coulouris G, Avagyan V, Ma N, Papadopoulos J, Bealer K, et al. BLAST+: architecture and applications. BMC bioinformatics. 2009;10(1):1–9.

37. Gee JE, Gulvik CA, Elrod MG, Batra D, Rowe LA, Sheth M, et al. Phylogeography of Burkholderia pseudomallei Isolates, Western Hemisphere. Emerg Infect Dis. 2017;23(7):1133–8.

38. Choi Y, Chan APJB. PROVEAN web server: a tool to predict the functional effect of amino acid substitutions and indels. 2015;31(16):2745–7.

39. Edgar RC. MUSCLE: multiple sequence alignment with high accuracy and high throughput. Nucleic Acids Res. 2004;32(5):1792–7.

40. Larkin MA, Blackshields G, Brown NP, Chenna R, McGettigan PA, McWilliam H, et al. Clustal W and Clustal X version 2.0. Bioinformatics. 2007;23(21):2947–8.

41. Cock PJ, Antao T, Chang JT, Chapman BA, Cox CJ, Dalke A, et al. Biopython: freely available Python tools for computational molecular biology and bioinformatics. Bioinformatics. 2009;25(11):1422–3.

42. Hunter PR, Gaston MA. Numerical index of the discriminatory ability of typing systems: an application of Simpson’s index of diversity. J Clin Microbiol. 1988;26(11):2465–6.

43. Nguyen LT, Schmidt HA, von Haeseler A, Minh BQ. IQ-TREE: a fast and effective stochastic algorithm for estimating maximum-likelihood phylogenies. Mol Biol Evol. 2015;32(1):268–74.

44. Letunic I, Bork PJNar. Interactive Tree Of Life (iTOL) v4: recent updates and new developments. 2019;47(W1):W256–W9.

45. Georgopapadakou NH, Liu FY. Penicillin-binding proteins in bacteria. Antimicrob Agents Chemother. 1980;18(1):148–57.

46. Krishnamurthy P, Parlow MH, Schneider J, Burroughs S, Wickland C, Vakil NB, et al. Identification of a novel penicillin-binding protein from Helicobacter pylori. J Bacteriol. 1999;181(16):5107–10.

47. Farra A, Islam S, Strålfors A, Sörberg M, Wretlind BJIjoaa. Role of outer membrane protein OprD and penicillin-binding proteins in resistance of Pseudomonas aeruginosa to imipenem and meropenem. 2008;31(5):427–33.

48. Kocaoglu O, Tsui HC, Winkler ME, Carlson EE. Profiling of beta-lactam selectivity for penicillin-binding proteins in Streptococcus pneumoniae D39. Antimicrob Agents Chemother. 2015;59(6):3548–55.

49. Sainsbury S, Bird L, Rao V, Shepherd SM, Stuart DI, Hunter WN, et al. Crystal structures of penicillin-binding protein 3 from Pseudomonas aeruginosa: comparison of native and antibiotic-bound forms. J Mol Biol. 2011;405(1):173–84.

50. Tomberg J, Temple B, Fedarovich A, Davies C, Nicholas RA. A highly conserved interaction involving the middle residue of the SXN active-site motif is crucial for function of class B penicillin-binding proteins: mutational and computational analysis of PBP 2 from N. gonorrhoeae. Biochemistry. 2012;51(13):2775–84.

51. Williamson R, Gutmann L, Horaud T, Delbos F, Acar JF. Use of penicillin-binding proteins for the identification of enterococci. J Gen Microbiol. 1986;132(7):1929–37.

52. McLaughlin HP, Bugrysheva J, Sue D. Optical microscopy reveals the dynamic nature of B. pseudomallei morphology during β-lactam antimicrobial susceptibility testing. J bioRxiv. 2020.

53. Lowe W, March, J. K., Bunnell, A. J., O’Neill, K. L., Robison, R. A. PCR-based Methodologies Used to Detect and Differentiate the Burkholderia pseudomallei complex: B. pseudomallei, B. mallei, and B. thailandensis. Curr Issues Mol Biol. 2013;16(2):23–54.

54. Kuhn G, Francioli P, Blanc D. Double-locus sequence typing using clfB and spa, a fast and simple method for epidemiological typing of methicillin-resistant Staphylococcus aureus. Journal of clinical microbiology. 2007;45(1):54–62.

55. Basset P, Blanc D. Fast and simple epidemiological typing of Pseudomonas aeruginosa using the double-locus sequence typing (DLST) method. European journal of clinical microbiology & infectious diseases. 2014;33(6):927–32.

56. Baker A, Pearson T, Price EP, Dale J, Keim P, Hornstra H, et al. Molecular phylogeny of Burkholderia pseudomallei from a remote region of Papua New Guinea. PLoS One. 2011;6(3):e18343.

57. Meumann EM, Kaestli M, Mayo M, Ward L, Rachlin A, Webb JR, et al. Emergence of Burkholderia pseudomallei Sequence Type 562, Northern Australia. Emerg Infect Dis. 2021;27(4):1057–67.

58. Chen H, Xia L, Zhu X, Li W, Du X, Wu D, et al. Burkholderia pseudomallei sequence type 562 in China and Australia. Emerging infectious diseases. 2015;21(1):166.

59. Birnie E, Virk HS, Savelkoel J, Spijker R, Bertherat E, Dance DA, et al. Global burden of melioidosis in 2015: a systematic review and data synthesis. The Lancet Infectious diseases. 2019;19(8):892–902.

