## Supplemental Tables for "*In silico* analyses of penicillin binding proteins in *Burkholderia pseudomallei* uncovers SNPs with utility for phylogeography, species differentiation, and sequence typing"

**Table S1**. Initial set of *Burkholderia* isolates analyzed with the corresponding genome accession numbers used.

| ***B. pseudomallei* isolate** | **Origin /**  **Patient Travel or Residence** | **Isolation Source** | **Isolation Year** | **GenBank Accessions** |
| --- | --- | --- | --- | --- |
| 1026b (Reference) | Thailand | Human blood (septicemic) | 1993 | CP002833, CP002834 |
| K96243 | Thailand | Human | 1993 | CP009537, CP009538 |
| BGR | Thailand | -- | -- | CP008834, CP008835 |
| PHLS112 | Thailand | Human clinical | 1992 | CP009585, CP009586 |
| 1710b | Thailand | -- | -- | CP000124, CP000125 |
| 406e | Thailand | Human | 1998 | CP009297, CP009298 |
| 1106a | Thailand | Pus (liver abscess) | 1993 | CP008758, CP008759 |
| Mahidol-1106a | Thailand | -- | -- | CP008781, CP008782 |
| 576 | Thailand | Human | -- | CP008777, CP008778 |
| FDAARGOS 592 | Thailand | Clinical | -- | CP033705, CP033706 |
| FDAARGOS 593 | -- | -- | -- | CP033703, CP033704 |
| FDAARGOS 594 | Thailand | -- | -- | CP033701, CP033702 |
| HBPUB10134a | Thailand | Human tracheal Suction | 2010 | CP008911, CP008912 |
| 14M0960418 | Hong Kong | Blood Culture | 2014 | CP019042, CP019043 |
| BPHN1 | China | Goat | 2016 | CP023775, CP023776 |
| 350105 | Hainin, China | Water | 1976 | CP012093, CP012094 |
| BPC006 | Hainin, China | Blood from TypeII diabetes patient with abscesses | 2008 | CP003781, CP003782 |
| vgh16W | Taiwan | Human blood | 2001 | CP012517, CP012518 |
| vgh16R | Taiwan | Human blood | 2001 | CP012515, CP012516 |
| vgh07 | Taiwan | Human blood | 1996 | CP010973, CP010974 |
| Pasteur 52237 | Vietnam | -- | -- | CP009898, CP009899 |
| MS | Malaysia | Human | 2015 | CP016636, CP016637 |
| PMC2000 | Malaysia | Human | 2000 | CP025302, CP025303 |
| D286 | Malaysia | Clinical | 1986 | CP025306, CP025307 |
| H10 | Pahang, Malaysia | Human | 1995 | CP025300, CP25301 |
| R15 | Malaysia | Human | 2005 | CP025304, CP025305 |
| 982 | Pahang, Malaysia | Pus | 2015 | CP012576, CP012577 |
| M1 | Malaysia | Human | 2015 | CP016638, CP016639 |
| Strain 9 | Pakistan | -- | -- | CP008754, CP008755 |
| VB3253 | India | Human blood aspirate | 2019 | CP040531, CP040532 |
| VB2514 | India | Human blood aspirate | 2019 | CP040551, CP040552 |
| Bps 110 | Sri Lanka | Human blood | 2015 | CP036451, CP036452 |
| Bps 111 | Sri Lanka | Human blood | 2015 | CP036453, CP036454 |
| Bps 112 | Sri Lanka | Human blood | 2015 | CP037975, CP037976 |
| Bps 114 | Sri Lanka | Human blood | 2015 | CP037973, CP037974 |
| Bps 115 | Sri Lanka | Human blood | 2015 | CP037757, CP037758 |
| Bps 116 | Sri Lanka | Human blood | 2015 | CP037759, CP037760 |
| Bps 122 | Sri Lanka | Human blood | 2015 | CP038194, CP038195 |
| Bps 123 | Sri Lanka | Human joint fluid/blood | 2015 | CP037969, CP037970 |
| Bps 133 | Sri Lanka | Human blood | 2015 | CP037971, CP037972 |
| BSR | -- | -- | -- | CP009127, CP009128 |
| BGK | Thailand | -- | -- | CP008916, CP008917 |
| NCTC 13178 | Australia | Human brain (*post mortem*) | -- | CP004001, CP004002 |
| NCTC 13179 | Australia | Human skin ulcer | -- | CP003976, CP003977 |
| NAU35A-3 | Australia | Soil | 2006 | CP004377, CP004378 |
| NAU20B-16 | Australia | Soil | 2006 | CP004003, CP004004 |
| TSV 202 | Australia | Environmental | -- | CP009156, CP009157 |
| TSV 48 | Australia | Environmental | -- | CP009160, CP009161 |
| BDP | Australia, NT | Brain | 1994 | CP009209, CP009210 |
| MSHR146 | Australia | Right udder (goat) | 1992 | CP004042, CP004043 |
| MSHR62 | Australia | Human clinical | -- | CP009235, CP009234 |
| MSHR5858 | Australia | Human sputum | 2011 | CP008891, CP008892 |
| MSHR2243 | Australia | Human clinical | -- | CP009270, CP009269 |
| MSHR840 | Australia, Ipswitch | Human brain tissue | 1999 | CP009473, CP009474 |
| MSHR511 | Australia | Throat (goat) | 1997 | CP004023, CP004024 |
| MSHR668 | Australia | Human brain | 1995 | CP009545, CP009546 |
| MSHR6755 | Australia, NT | Blood | 2012 | CP017046, CP017047 |
| MSHR4083 | Australia, NT | Human | 2010 | CP017050, CP017051 |
| MSHR7929 | Australia, NT | Blood | 2013 | CP017044, CP017045 |
| MSHR520 | Australia | Blood | 1998 | CP004368, CP004369 |
| MSHR305 | Australia, NT | Brain sample (autopsy) | 1994 | CP006469, CP006470 |
| MSHR5864 | Australia, NT | Blood | 2011 | CP017048, CP017049 |
| MSHR3763 | Australia, NT | Human | 2010 | CP017052, CP017053 |
| MSHR2543 | Australia | -- | -- | CP009477, CP009478 |
| MSHR491 | Australia, NT | Community water supply storage tank | 1997 | CP009484, CP009485 |
| MSHR1153 | Australia | Clinical | -- | CP009271, CP009272 |
| MSHR1435 | Australia, NT | Environment | 2002 | CP025264, CP025265 |
| MSHR3965 | Australia | Environment | -- | CP009152, CP009153 |
| MSHR1655 | Australia | -- | -- | CP008779, CP008780 |
| Bp1651 | USA / Australia | Human sputum | -- | CP012041, CP012042 |
| Burk178-Type1 | Australia | Human sputum | 2011 | CP016909, CP016910 |
| Burk179-Type2 | Australia | Human sputum | 2011 | CP016911, CP016912 |
| K42 | Papua New Guinea | Environmental | -- | CP009162, CP00963 |
| B03 | Papua New Guinea | Environmental | -- | CP009150, CP009151 |
| A79A | Papua New Guinea | Environmental | -- | CP009165, CP009164 |
| VB976100 | Czech Republic | Pus of abscess (iguana) | 2014 | CP018054, CP018055 |
| PR1998 | Puerto Rico | Human clinical | 1998 | CP018369, CP018370 |
| PR1982 | Puerto Rico | Human clinical | 1982 | CP018367, CP018368 |
| PR2012 | Puerto Rico | Human clinical | 2012 | CP018393, CP018394 |
| PR2013a | Puerto Rico | Soil | 2013 | CP018406, CP018407 |
| PR2013b | Puerto Rico | Soil | 2103 | CP018408, CP018409 |
| FL2012 | FL, USA / Trinidad | Human clinical | 2012 | CP018391, CP018392 |
| MX2013 | CA, USA /  Mexico & Vietnam | Human clinical | 2013 | CP018395, CP018396, CP018397 |
| TX2004 | TX, USA / SE Asia | Human clinical | 2004 | CP018375, CP018376 |
| IL2014 | IL, USA / Mexico | Human clinical | 2014 | CP018414, CP018415 |
| VEN1976 | Venezuela / Unknown | Human clinical | 1976 | CP018371, CP018372 |
| 7894 | Ecuador / Unknown | Human clinical | 1962 | CP018373, CP018374 |
| CA2007 | CA, USA / Unknown | Human clinical | 2007 | CP018418, CP018419 |
| CA2009 | CA, USA / Mexico | Human clinical | 2009 | CP018380, CP018381 |
| PB1007001 | AZ, USA / Costa Rica | Human clinical | 2009 | CP018387, CP018388 |
| OH2013 | OH, USA / None | Human clinical | 2013 | CP018400, CP018401 |
| NY2010 | NY, USA / Aruba | Human clinical | 2010 | CP018384, CP018386 |
| Swiss2010 | Switzerland / Martinique | Human clinical | 2010 | CP018389, CP018390 |
| GA2015 | GA, USA / Panama & Peru | Human clinical | 2015 | CP018416, CP018417 |
| TX2015 | TX, USA / Mexico | Human clinical | 2015 | CP018412, CP018413 |
| CA2010 | CA, USA / Unknown | Human clinical | 2010 | CP018382, CP018383 |
| CA2013a | CA, USA / Unknown | Human clinical | 2013 | CP018398, CP018399 |
| MX2014 | CA, USA / Mexico | Human clinical | 2014 | CP018410, CP018411 |
| RI2013a | RI, USA / Guatemala | Human clinical | 2013 | CP018402, CP018403 |
| RI2013b | RI, USA / Guatemala | Human clinical | 2013 | CP018404, CP018405 |
| PB08298010 | AZ, USA / Unknown | Human clinical | 2008 | CP009550, CP009551 |
| ***B. mallei* isolate** | **Origin** | **Source** | **Year** | **Accession** |
| ATCC23344 | -- | -- | -- | CP000010, CP000011 |
| NCTC10247 | Turkey | -- | -- | CP007801, CP007802 |
| Bahrain1 | Bahrain | Horse | 2011 | CP017175, CP017176 |
| JHU | USA | Human | 2000 | CP009931, CP009932 |
| FMH | USA | Human | 2000 | CP009929, CP009930 |
| 2002721276 | -- | -- | 1956 | CP010065, CP010066 |
| 2002734306 | UK | -- | -- | CP009707, CP009708 |
| India86-567-2 | India | Mule |  | CP009642, CP009643 |
| strain 11 | Turkey | Human | 1949 | CP009587, CP009588 |
| strain 6 | -- | -- | -- | CP008710, CP008711 |
| 2002734299 | Hungary | Guinea Pig | 1961 | CP009337, CP009338 |
| BMQ | India | Horse |  | CP008722, CP008723 |
| NCTC10229 | -- | -- | -- | CP000545, CP000546 |
| SAVP1 | -- | -- | -- | CP000525, CP000526 |
| KC_1092 | Iran | -- | -- | CP009942, CP009943 |
| 2000031063 | -- | -- | -- | CP008731, CP008732 |
| Turkey10 | Turkey | -- | -- | CP010348, CP010349 |
| Turkey9 | Turkey | -- | -- | CP009741, CP009742 |
| Turkey8 | Turkey | -- | -- | CP009739, CP009740 |
| Turkey7 | Turkey | -- | -- | CP009737, CP009738 |
| Turkey6 | Turkey | -- | -- | CP009735, CP009736 |
| Turkey5 | Turkey | -- | -- | CP009733, CP009734 |
| Turkey4 | Turkey | -- | -- | CP009731, CP009732 |
| Turkey3 | Turkey | -- | -- | CP009729, CP009730 |
| Turkey2 | Turkey | -- | -- | CP009727, CP009728 |
| Turkey1 | Turkey | -- | -- | CP009725, CP009726 |
| ***B. thailandensis* isolate** | **Origin** | **Source** | **Year** | **Accession** |
| E444 | Thailand | Soil | 2002 | CP004117, CP004118 |
| E254 | Thailand | -- | 1992 | CP004381, CP004382 |
| E264 | Thailand | Rice Field Soil | -- | CP008785, CP008786 |
| H0587 | LA, USA | Human Pleural Wound | 1997 | CP004089 CP004090 |
| MSMB59 | Australia, NT | Soil | 2006 | CP013407 CP013408 |
| FDAARGOS 237 | USAMRIID | Water | 2009 | CP020390, CP020389 |
| FDAARGOS 238 | USAMRIID | Human Pleural Wound | 2009 | CP020392, CP020391 |
| FDAARGOS 241 | USAMRIID/CDC | -- | 2011 | CP022214, CP022215 |
| FDAARGOS 242 | Thailand | -- | 2011 | CP022217 CP022216 |
| FDAARGOS 426 | USAMRIID | Environment | -- | CP023499, CP023498 |
| MSMB121 | Australia | Soil | 2007 | CP004095, CP004096 |
| 20027211643 | -- | -- | 2002 | CP009601, CP009602 |
| 34 | -- | -- | 2002 | CP010017, CP010018 |
| USAMRU Malaysia #20 | Malaysia | -- | -- | CP004383, CP004384 |
| 2002721121 | USA | -- | -- | CP013409 CP013410 |
| 2003015869 | TX, USA | Human NAU subculture | 2003 | CP013360, CP013361 |
| 2002721723 | -- | Human | -- | CP004097, CP004098 |

Summary of epidemiological data and assembled genome sequences publicly available (GenBank, NCBI) for the initial set of *B. pseudomallei* (101), *B. mallei* (26) and *B*. *thailandensis* (17) strains used for SNP analyses of PBPs. Information not available (--), Australia, Northern Territory (NT).

**Table S2**. Dual Locus Sequence Typing (DLST) results for the initial set of *B. pseudomallei* and *B. mallei* strains.

| ***B. pseudomallei* isolate** | **DLST** | **nucleotide position (*I0276*)** | | | | | **nucleotide position (*II1314)*** | | | | | |
| --- | --- | --- | --- | --- | --- | --- | --- | --- | --- | --- | --- | --- |
|  | **Sequence Type** | **141** | **268** | **888** | **1473** | **1629** | **243** | **265** | **273** | **575** | **703** | **854** |
| 1026b (Reference) | 1 | C | C | C | C | T | G | G | T | T | A | A |
| K96243 | 1 | C | C | C | C | T | G | G | T | T | A | A |
| BGR | 1 | C | C | C | C | T | G | G | T | T | A | A |
| PHLS112 | 5 | C | C | C | C | T | A | G | C | C | A | C |
| 1710b | 1 | C | C | C | C | T | G | G | T | T | A | A |
| 406e | 13 | C | C | C | C | T | G | G | C | C | A | A |
| 1106a | 4 | C | C | C | C | T | G | G | C | T | A | A |
| Mahidol-1106a | 4 | C | C | C | C | T | G | G | C | T | A | A |
| 576 | 3 | C | C | C | C | T | G | G | C | C | A | C |
| FDAARGOS 592 | 3 | C | C | C | C | T | G | G | C | C | A | C |
| FDAARGOS 593 | 5 | C | C | C | C | T | A | G | C | C | A | C |
| FDAARGOS 594 | 3 | C | C | C | C | T | G | G | C | C | A | C |
| HBPUB10134a | 1 | C | C | C | C | T | G | G | T | T | A | A |
| 14M0960418 | 4 | C | C | C | C | T | G | G | C | T | A | A |
| BPHN1 | 13 | C | C | C | C | T | G | G | C | C | A | A |
| 350105 | 24 | C | C | C | C | T | G | G | C | C | G | A |
| BPC006 | 4 | C | C | C | C | T | G | G | C | T | A | A |
| vgh16W | 1 | C | C | C | C | T | G | G | T | T | A | A |
| vgh16R | 1 | C | C | C | C | T | G | G | T | T | A | A |
| vgh07 | 1 | C | C | C | C | T | G | G | T | T | A | A |
| Pasteur 52237 | 3 | C | C | C | C | T | G | G | C | C | A | C |
| MS | 9 | C | T | C | C | T | G | G | C | C | G | C |
| PMC2000 | 4 | C | C | C | C | T | G | G | C | T | A | A |
| D286 | 4 | C | C | C | C | T | G | G | C | T | A | A |
| H10 | 1 | C | C | C | C | T | G | G | T | T | A | A |
| R15 | 4 | C | C | C | C | T | G | G | C | T | A | A |
| 982 | 1 | C | C | C | C | T | G | G | T | T | A | A |
| M1 | 9 | C | T | C | C | T | G | G | C | C | G | C |
| Strain 9 | 3 | C | C | C | C | T | G | G | C | C | A | C |
| VB3253 | 6 | C | C | C | C | T | A | G | C | C | G | C |
| VB2514 | 6 | C | C | C | C | T | A | G | C | C | G | C |
| Bps 110 | 2 | C | C | C | C | T | G | G | C | C | G | C |
| Bps 111 | 2 | C | C | C | C | T | G | G | C | C | G | C |
| Bps 112 | 6 | C | C | C | C | T | A | G | C | C | G | C |
| Bps 114 | 6 | C | C | C | C | T | A | G | C | C | G | C |
| Bps 115 | 6 | C | C | C | C | T | A | G | C | C | G | C |
| Bps 116 | 6 | C | C | C | C | T | A | G | C | C | G | C |
| Bps 122 | 6 | C | C | C | C | T | A | G | C | C | G | C |
| Bps 123 | 2 | C | C | C | C | T | G | G | C | C | G | C |
| Bps 133 | 6 | C | C | C | C | T | A | G | C | C | G | C |
| BSR | 13 | C | C | C | C | T | G | G | C | C | A | A |
| BGK | 1 | C | C | C | C | T | G | G | T | T | A | A |
| NCTC 13178 | 2 | C | C | C | C | T | G | G | C | C | G | C |
| NCTC 13179 | 2 | C | C | C | C | T | G | G | C | C | G | C |
| NAU35A-3 | 7 | C | C | C | C | T | G | G | C | T | G | C |
| NAU20B-16 | 2 | C | C | C | C | T | G | G | C | C | G | C |
| TSV 202 | 3 | C | C | C | C | T | G | G | C | C | A | C |
| TSV 48 | 2 | C | C | C | C | T | G | G | C | C | G | C |
| BDP | 7 | C | C | C | C | T | G | G | C | T | G | C |
| MSHR146 | 2 | C | C | C | C | T | G | G | C | C | G | C |
| MSHR62 | 2 | C | C | C | C | T | G | G | C | C | G | C |
| MSHR5858 | 24 | C | C | C | C | T | G | G | C | C | G | A |
| MSHR2243 | 2 | C | C | C | C | T | G | G | C | C | G | C |
| MSHR840 | 3 | C | C | C | C | T | G | G | C | C | A | C |
| MSHR511 | 2 | C | C | C | C | T | G | G | C | C | G | C |
| MSHR668 | 2 | C | C | C | C | T | G | G | C | C | G | C |
| MSHR6755 | 2 | C | C | C | C | T | G | G | C | C | G | C |
| MSHR4083 | 7 | C | C | C | C | T | G | G | C | T | G | C |
| MSHR7929 | 2 | C | C | C | C | T | G | G | C | C | G | C |
| MSHR520 | 7 | C | C | C | C | T | G | G | C | T | G | C |
| MSHR305 | 7 | C | C | C | C | T | G | G | C | T | G | C |
| MSHR5864 | 2 | C | C | C | C | T | G | G | C | C | G | C |
| MSHR3763 | 7 | C | C | C | C | T | G | G | C | T | G | C |
| MSHR2543 | 9 | C | T | C | C | T | G | G | C | C | G | C |
| MSHR491 | 2 | C | C | C | C | T | G | G | C | C | G | C |
| MSHR1153 | 2 | C | C | C | C | T | G | G | C | C | G | C |
| MSHR1435 | 2 | C | C | C | C | T | G | G | C | C | G | C |
| MSHR3965 | 2 | C | C | C | C | T | G | G | C | C | G | C |
| MSHR1655 | 2 | C | C | C | C | T | G | G | C | C | G | C |
| Bp1651 | 2 | C | C | C | C | T | G | G | C | C | G | C |
| Burk178-Type1 | 7 | C | C | C | C | T | G | G | C | T | G | C |
| Burk179-Type2 | 7 | C | C | C | C | T | G | G | C | T | G | C |
| K42 | 18 | C | C | C | T | T | G | G | C | C | G | C |
| B03 | 18 | C | C | C | T | T | G | G | C | C | G | C |
| A79A | 18 | C | C | C | T | T | G | G | C | C | G | C |
| VB976100 | 17 | T | C | C | C | T | G | G | C | T | A | A |
| PR1998 | 12 | C | T | C | C | T | G | T | T | T | A | A |
| PR1982 | 12 | C | T | C | C | T | G | T | T | T | A | A |
| PR2012 | 12 | C | T | C | C | T | G | T | T | T | A | A |
| PR2013a | 12 | C | T | C | C | T | G | T | T | T | A | A |
| PR2013b | 12 | C | T | C | C | T | G | T | T | T | A | A |
| FL2012 | 12 | C | T | C | C | T | G | T | T | T | A | A |
| MX2013 | 1 | C | C | C | C | T | G | G | T | T | A | A |
| TX2004 | 10 | C | T | C | C | T | G | G | T | T | A | A |
| IL2014 | 10 | C | T | C | C | T | G | G | T | T | A | A |
| VEN1976 | 10 | C | T | C | C | T | G | G | T | T | A | A |
| 7894 | 10 | C | T | C | C | T | G | G | T | T | A | A |
| CA2007 | 25 | C | T | C | C | T | G | G | T | C | A | A |
| CA2009 | 11 | C | T | C | C | T | G | G | C | C | A | A |
| PB1007001 | 10 | C | T | C | C | T | G | G | T | T | A | A |
| OH2013 | 3 | C | C | C | C | T | G | G | C | C | A | C |
| NY2010 | 10 | C | T | C | C | T | G | G | T | T | A | A |
| Swiss2010 | 10 | C | T | C | C | T | G | G | T | T | A | A |
| GA2015 | 17 | T | C | C | C | T | G | G | C | T | A | A |
| TX2015 | 22 | T | C | C | C | T | G | G | C | C | A | A |
| CA2010 | 5 | C | C | C | C | T | A | G | C | C | A | C |
| CA2013a | 25 | C | T | C | C | T | G | G | T | C | A | A |
| MX2014 | 17 | T | C | C | C | T | G | G | C | T | A | A |
| RI2013a | 17 | T | C | C | C | T | G | G | C | T | A | A |
| RI2013b | 17 | T | C | C | C | T | G | G | C | T | A | A |
| PB08298010 | 3 | C | C | C | C | T | G | G | C | C | A | C |
| ***B. mallei* isolate** | **DLST** | **nucleotide position (*I0276*)** | | | | | **nucleotide position (*II1314)*** | | | | | |
|  | **Sequence Type** | **141** | **268** | **888** | **1473** | **1629** | **243** | **265** | **273** | **575** | **703** | **854** |
| ATCC23344 | 32 | C | T | T | C | C | A | G | C | C | G | C |
| NCTC10247 | 32 | C | T | T | C | C | A | G | C | C | G | C |
| Bahrain1 | 32 | C | T | T | C | C | A | G | C | C | G | C |
| JHU | 32 | C | T | T | C | C | A | G | C | C | G | C |
| FMH | 32 | C | T | T | C | C | A | G | C | C | G | C |
| 2002721276 | 32 | C | T | T | C | C | A | G | C | C | G | C |
| 2002734306 | 32 | C | T | T | C | C | A | G | C | C | G | C |
| India86-567-2 | 32 | C | T | T | C | C | A | G | C | C | G | C |
| strain 11 | 32 | C | T | T | C | C | A | G | C | C | G | C |
| strain 6 | 32 | C | T | T | C | C | A | G | C | C | G | C |
| 2002734299 | 32 | C | T | T | C | C | A | G | C | C | G | C |
| BMQ | 32 | C | T | T | C | C | A | G | C | C | G | C |
| NCTC10229 | 32 | C | T | T | C | C | A | G | C | C | G | C |
| SAVP1 | 32 | C | T | T | C | C | A | G | C | C | G | C |
| KC_1092 | 32 | C | T | T | C | C | A | G | C | C | G | C |
| 2000031063 | 32 | C | T | T | C | C | A | G | C | C | G | C |
| Turkey10 | 32 | C | T | T | C | C | A | G | C | C | G | C |
| Turkey9 | 32 | C | T | T | C | C | A | G | C | C | G | C |
| Turkey8 | 32 | C | T | T | C | C | A | G | C | C | G | C |
| Turkey7 | 32 | C | T | T | C | C | A | G | C | C | G | C |
| Turkey6 | 32 | C | T | T | C | C | A | G | C | C | G | C |
| Turkey5 | 32 | C | T | T | C | C | A | G | C | C | G | C |
| Turkey4 | 32 | C | T | T | C | C | A | G | C | C | G | C |
| Turkey3 | 32 | C | T | T | C | C | A | G | C | C | G | C |
| Turkey2 | 32 | C | T | T | C | C | A | G | C | C | G | C |
| Turkey1 | 32 | C | T | T | C | C | A | G | C | C | G | C |

DLST typing results for the initial set of *Burkholderia* strains analyzed and reported in Figure 1. Nucleotides with phylogeographic utility are shown in black font and nucleotides used to differentiate *Burkholderia* species are shown in gray font.

**Table S3**. DLST results and geographic origin for the 1,523 *B. pseudomallei* and *B. mallei* RefSeq genomes.

| **ST** | **DLST Sequence** | **Total # of Genomes** | **Country (# of strains)** |
| --- | --- | --- | --- |
| 1 | CCCCTGGTTAA | 451 | Thailand (416), Singapore (11), Malaysia (8), Vietnam (5), UK (4)^+^, Taiwan (3), Bangladesh (1), China (1), New Zealand (1), USA (1) |
| 2 | CCCCTGGCCGC | 220 | Australia (110), Thailand (100), Malaysia (5), Sri Lanka (3), USA (2)^#^ |
| 3 | CCCCTGGCCAC | 205 | Thailand (105), Singapore (48), Australia (30), Malaysia (7), France (5), Pakistan (3), Bangladesh (2), Papua New Guinea (1), Portugal (1), UK (1), USA (1), Vietnam (1) |
| 4 | CCCCTGGCTAA | 157 | Thailand (122), Singapore (26), Malaysia (5), China (1), Hong Kong (1), South Korea (1), Vietnam (1) |
| 5 | CCCCTAGCCAC | 92 | Thailand (91), USA (1)* |
| 6 | CCCCTAGCCGC | 74 | Thailand (66), Sri Lanka (6), Israel (1), Vietnam (1) |
| 7 | CCCCTGGCTGC | 67 | Australia (63), Thailand (4) |
| 8 | CTCCTAGCCAC | 34 | Thailand (34) |
| 9 | CTCCTGGCCGC | 32 | Australia (23), Malaysia (8), Thailand (1) |
| 10 | CTCCTGGTTAA | 13 | USA (9), Ecuador (2), Switzerland (1) Venezuela (1) |
| 11 | CTCCTGGCCAA | 13 | Australia (12), USA (1) |
| 12 | CTCCTGTTTAA | 11 | USA (6), Puerto Rico (5) |
| 13 | CCCCTGGCCAA | 10 | Thailand (7), Australia (2), China (1) |
| 14 | CTCCTGGCCAC | 8 | Australia (6), New Zealand (1), Thailand (1) |
| 15 | CCCCTGGTTAC | 7 | Thailand (7) |
| 16 | CCCCTAGCCAA | 6 | Thailand (6) |
| 17 | TCCCTGGCTAA | 5 | USA (4), Czech Republic (1) |
| 18 | CCCTTGGCCGC | 5 | Papua New Guinea (5) |
| 19 | CCCCTGGTCAC | 5 | Thailand (5) |
| 20 | CCCCTGGCTAC | 5 | Australia (4), Thailand (1) |
| 21 | CCCCTAGCTGC | 4 | Thailand (4) |
| 22 | TCCCTGGCCAA | 3 | Mexico (2), USA (1) |
| 23 | CCCCTGGTCAA | 3 | Thailand (3) |
| 24 | CCCCTGGCCGA | 3 | Australia (3) |
| 25 | CTCCTGGTCAA | 2 | USA (2) |
| 26 | CCCCTGGTTGC | 2 | Thailand (2) |
| 27 | CTCCTGGCTGC | 1 | Australia (1) |
| 28 | CCCCTGGTCGC | 1 | Thailand (1) |
| 29 | CCCCTGGACGC | 1 | Australia (1) |
| 30 | CCCCTGGACAC | 1 | Australia (1) |
| 31 | CCCCTAGCCGA | 1 | Thailand (1) |
| 32 | CTTCCAGCCGC  (*B. mallei*) | 81 | 55/81 strains have associated geographic information:  Turkey (24), India (8), USA (6), Hungary (5), China (4), Myanmar (2), UK (2), Bahrain (1), Iran (1), Pakistan (1), Russia (1) |

Sequence typing results and epidemiological data for 1,523 *Burkholderia* strains used to assess the performance of the DLST approach. All RefSeq genomes are publicly available (NCBI) and geographic information was acquired from the BioSample database. DLSTs were assigned in numerical order based on the number of strains in each ST, highest to lowest. (^+^) UK laboratory cultures of *B. pseudomallei* K96243, originally from Thailand (Wagley *et al*, 2019). (*) ITS Type C. (^#^) 1 of 2 strains is Bp1651, Australian in origin.
